# Drug tolerant persisters and immunotherapy persister cells exhibit cross-resistance and share common survival mechanisms

**DOI:** 10.1101/2025.03.04.641492

**Authors:** Maria Davern, Cole J Turner, Daryl Griffin, Lara Bencsics, Brenda C Chan, Jasmine Yun-Tong Kung, Michael L Olson, Cheyanne Walker Williams, Shaili Soni, Leah Krotee, Michael Yorsz, Gabriella Antonellis, Patrick H Lizotte, Cloud P Paweletz, Jeremy Ryan, Filippo Birocchi, Antonio Josue Almazan, Kristopher A Sarosiek, David Barbie, Patrick Bhola, Marcela V Maus, Anthony Letai

## Abstract

Persisters are a rare sub-population of tumor cells that survive anti-cancer therapy and are thought to be a major cause of recurrence. These cells have been identified following both drug- and immune-therapy but are generally considered to be distinct entities. Since both pharmacological agents and immune cells often kill via apoptosis, we tested a hypothesis that both types of cells survive based on reduced mitochondrial apoptotic sensitivity, which in turn would yield a similar and reciprocal multi-agent resistant phenotype. Supporting this hypothesis, we indeed observed that IPCs acquired a reduced sensitivity to multiple drug classes and radiotherapy, suggesting non-immune mechanisms are important in the survival of cancer cells after immunotherapy. Likewise, DTPs developed not only a reduced sensitivity to multiple drug classes and radiotherapy, but also acquired a reduced sensitivity to T cell killing. Both IPCs and DTPs developed a decreased sensitivity to mitochondrial apoptosis. A sub-population of IPCs downregulated antigen and upregulated PD-L1. Intriguingly, in the IPCs that didn’t employ these mechanisms of resistance, a greater decrease in sensitivity to mitochondrial apoptosis was observed, suggesting that the presence or absence of a resistance mechanism can exert selective pressures over the emergence of others. Targeting anti-apoptotic dependencies in persisters increased sensitivity to chemotherapy or CAR T therapy. These results suggest that common biological mechanisms underly survival of persisters, whether derived from immune or drug therapy, and offer an explanation for the acquired cross-resistance to these two types of therapies often observed in the clinic.

**Highlights:**

- Immunotherapy persister cells (IPCs) are less sensitive to drugs and radiation.
- Drug tolerant persisters (DTPs) are less sensitive to radiation and CAR T cell attack.
- IPCs and DTPs are less sensitive to mitochondrial apoptosis.
- Targeting anti-apoptotic dependencies helps eliminate IPCs/DTPs.

**Graphical Abstract:** 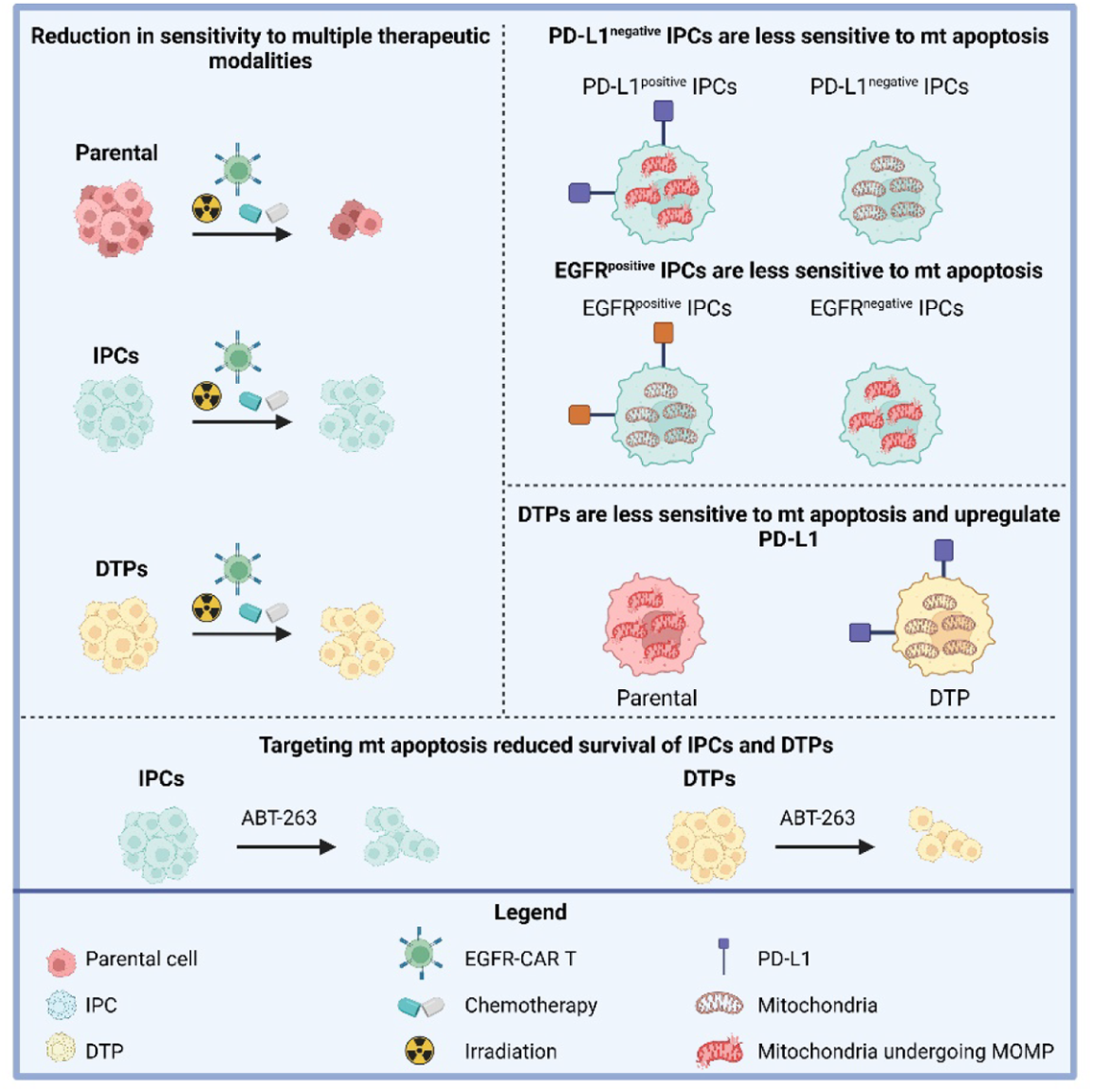

## Introduction

Persister tumor cells survive cancer therapy, are typically undetectable by PET/CT scans and are thought to give rise to tumor recurrence^1^. These persister cancer cells constitute a major obstacle for successful cancer treatment and are thought to be a key cause of tumor relapses that can occur months or years after initially successful cancer treatment^2^. These cells have been described as “that army known as invincible in peace, invisible in war”^3^. They reside in a slow-dividing cell state, and in response to signals that are not fully understood they can reawaken and resume growing again resulting in tumor recurrence and metastasis^4^. This has made complete tumor eradication very difficult to achieve and highlights the unmet clinical need to identify drugs that specifically target these persister cells that remain after treatment. The majority of what we have learned about persister cells has been achieved by studying tumor cells that persist following drug therapy. Considerable strides have been made in understanding drug tolerant persister (DTPs) cells which are characterized by their slow proliferation, reduction in sensitivity to the drug they survived and phenotypic plasticity, in that they can re-enter the cell cycle and regrow a tumor^5^. Epigenetic processes as opposed to genetic mechanisms have been identified to regulate the development of DTP tumor cells^5^.

Relapse and resistance following immune therapy is also a major clinical problem in oncology. Most studies of acquired resistance to immune therapy are concerned with the immune environment or the interaction of target tumor cells with immune cells^6,7^. There is relatively less consideration given to the response determinants that are purely target-cell intrinsic. However, persister cells have also been identified following immunotherapy treatment and it remains unknown how different from or similar to each other these persister cell fates are. Sehgal et al^8^, identified a cancer cell subpopulation expressing stem-like markers Snail and Sca-1 that had escaped CD8^+^ T cell-mediated killing after effective PD-1 blockade, which they called immunotherapy persister cells (IPCs). The authors successfully enhanced response to PD-1 blockade in mice by therapeutically targeting an apoptotic pathway in tumor cells using a SMAC mimetic that targeted Birc2/3 in combination with anti-PD-1 therapy^8^. This combination resulted in durable responses by decreasing the surviving fraction of IPCs^8^. A complementary study by Joung et al^9^, suggests a potential role for alterations in mitochondrial apoptotic pathways in tumor cells as a mechanism to survive T cell attack. Using a CRISPR activation screen, Joung et al^9^, demonstrated that tumor cell resistance to T cell-mediated killing was driven by alterations in mitochondrial apoptotic BCL-2 proteins and B3GNT2. Collectively, these two studies implicate a potential role for alterations in mitochondrial apoptotic pathways in providing tumor cells with a survival advantage against T cell killing. This highlights a promising exploitable therapeutic approach utilizing drugs that target the mitochondrial apoptotic pathway to improve T cell killing of target tumor cells and the subsequent efficacy of immune checkpoint blockade or CAR T cell therapy. Another study highlighted the importance of the apoptotic pathway for T cell-mediated killing of tumor cells where they showed that granzyme derived from the cytotoxic granules of T cells activated the mitochondrial apoptotic cell death pathway in target cells^10^.

The objective of this study is to ask if there are universal properties common among persister cells emerging from both drug and immune cell therapy. A diverse array of resistance mechanisms selective against different classes of chemotherapies have been reported which can be regulated by genetic or epigenetic mechanisms^11,12^. We hypothesize that persister tumor cells could have a shared mechanism of resistance that might be present across different persister cells independent of the drug or cellular therapy used to generate the persister cells. Therefore, tumor cells might employ common pathways to cope with cellular stress irrespective of the therapeutic insult, which could promote a shared mechanism of survival common to many persister cells. Considering that many drugs^13^ and T cells kill tumor cells via mitochondrial apoptosis^14^, we propose that alterations in mitochondrial apoptotic signaling could be a shared mechanism of resistance across DTPs and IPCs. Hence, in this study we sought to investigate if DTPs and IPCs are less sensitive (less “primed”) for mitochondrial apoptosis. We utilize a functional assay known as BH3 profiling^15^ to assess how close a cell is to the “threshold” of mitochondrial apoptosis^15^. An important implication of this hypothesis is that one might then expect persisters derived from immune therapy to be resistant not only to immune attack, but also to drug treatment. Conversely, persisters derived from drug treatment would be expected to be resistant not only to drug treatment, but also to immune therapy.

Use of drugs to target mitochondrial apoptotic signaling in tumor cells has proven very useful in improving the efficacy of many chemotherapies^15–18^ as well as immunotherapies such as PD-1 immune checkpoint blockade^19^, NK cell therapy^20^ and CAR T cell therapy^21^. Therefore, in our study we employ BH3 profiling^15^ to also probe for anti-apoptotic dependencies in persisters to guide the selection of the “right” BH3 mimetic to increase the sensitivity of persisters to mitochondrial apoptotic cell death and improve their clearance.

## Materials and Methods

### Cell lines

HeLa and U251 human tumor cell lines were obtained from ATCC (American Type Culture Collection) and cultured in Dulbecco’s Modified Eagle Medium (DMEM) (Life Technologies, 11995073) supplemented with 10% fetal bovine serum (FBS) (Life Technologies, 10437028) and 1% penicillin-streptomycin (Life Technologies, 15140122). B16-OVA and MC38-OVA tumor cell lines were kindly gifted by Dr. Patrick Lizotte from Cloud Paweletz’s lab in Dana-Farber Cancer Institute, Boston, USA. Cell lines were regularly tested for mycoplasma contamination.

### Drug tolerant persister cell (DTP) generation

HeLa tumor cell lines were seeded in vented T300 cm^3^ flasks each at a density of 2.5 x 10^6^ cells per flask in 35 ml of media. B16-OVA and MC38-OVA tumor cell lines were seeded in vented T300 cm^3^ flasks each at a density of 3 x 10^6^ cells per flask in 35 ml of media. Cells were left to adhere overnight at 37°C, 5% CO_2_. The next day, cells were treated with an IC_90_ dose of drug (a dose that killed 90% of the cells within 72h) for 7 days to generate persister cells. These 72h IC_90_ doses were determined using a CellTiter-Glo assay (Promega, G7573). The drugged media was replenished every 3 days with freshly drugged media that contained an IC_90_ dose of the drug. The drugs used to generate persister cells included: 5-fluoruracil (5-FU) (Millipore Sigma, F6627), oxaliplatin (Selleckchem, S1224), etoposide (Selleckchem, S1225). Following a 7-day drug treatment, the flasks containing the DTPs were washed with phosphate buffered saline (PBS) (Life Technologies, 10010049) and trypsinized with 5 ml of 10X trypsin (Millipore Sigma, T4174) for 5 mins at 37°C, 5% CO_2_ and neutralized with 10 ml of media. Cells were then counted using trypan blue to exclude dead cells on the Countess 3FL (Invitrogen, Thermo Fisher Scientific) and used for further experimentation.

### EGFR-CAR T cell generation

T cells were isolated from normal donor PBMCs and lentivirally transduced with second generation chimeric antigen receptors (CARs) targeting EGFR with a 4-1BB co-stimulatory domain, similar to previously published protocols by the Maus Laboratory^22^.

### Mice

8-12-week-old-female C57/BL6 mice (The Jackson Laboratory, RRID:IMSR_JAX:000058) were used in all experiments. All mice were housed in pathogen-free facilities, in a 12-hour light/dark cycle in ventilated cages, with chow and water supply ad libitum.

### Ethics

All mice were maintained within the DCFI animal facility. All experiments involving animals received full ethical approval and were conducted in accordance with the DFCI policy and animal protocol, reviewed and approved by the DFCI Institutional Animal Care and Use Committee (IACUC approved protocol #18–005).

### Murine spleen digestion, CD3^+^ T cell isolation and activation

Spleens were removed from 8–12-week-old female C57BL/6 mice (The Jackson Laboratory, RRID:IMSR_JAX:000058). Each spleen was mechanically digested in a falcon 100 µM cell strainer (Life Sciences, 352360) using the back of a 2 ml plunger and the cells were washed through the filter using PBS (Life Technologies, 10010049) containing 2% FBS (Life Technologies, 10437028) and 1 mM EDTA (ThermoFisher Scientific, 15575020). From the splenocyte cell suspension, CD3^+^ T cells were isolated using the EasySep™ Mouse CD3^+^ T Cell Isolation Kit following the manufacturer’s instructions. CD3^+^ T cells were activated overnight using Dynabeads® Mouse T-Activator CD3/CD28 (Life Technologies, 11452D) in a bead-to-cell ratio of 1:1 in Roswell Parks Memorial Institute (RPMI)-1640 (Invitrogen, 7410) supplemented with 10% fetal bovine serum (FBS) (Life Technologies, 10437028) and 1% penicillin-streptomycin (Life Technologies, 15140122) and IL-2 (30 IU/ml) (Peprotech, 200-02). Activation beads were removed and activated T cells were then used in experiments.

### Immunotherapy persister cell (IPC) generation

HeLa and U251 tumor cell lines were seeded in 12 cm^3^ petri dishes at a density of 0.5 x 10^6^ cells per dish in 10 ml of media. Cells were left to adhere overnight at 37°C, 5% CO_2_. The next day, the media was refreshed and EGFFR-CAR T cells were added at an effector to target E:T ratio of 0.2:1 or 0.3:1 for Hela and U251 cell lines, respectively. Every 3 days the media was replaced and fresh CAR T cells were added at the same density. The co-culture was maintained for up to 3-4 wks. At the end of the co-culture, the plates containing the IPCs were washed with PBS (Life Technologies, 10010049) and trypsinized with 1.5 ml of trypsin for 5 mins at 37°C, 5% CO_2_. Cells were then counted using trypan blue to exclude dead cells on the Countess 3FL (Invitrogen, Thermo Fisher Scientific) and used for further experimentation.

### Microscopy-based BH3 profiling and dynamic BH3 profiling

BH3 profiling and dynamic BH3 profiling (DBP) were carried on cell lines seeded at a density of 5 x 10^3^ cells/well in 30 ul of media in a 384-well plate. For BH3 profiling the cells were not drugged and for DBP cells were drug treated 1 hour after cell seeding using the D300e digital drug dispenser (Hewlett Packard). The next day the media/drugs were washed from plates using the BioTek 406EL plate washer (BioTek) and replaced with 20 µl of PBS per well. BH3 profiling buffer at a concentration of 2X (1× is 150 nM Mannitol, 10 mM HEPES-KOH pH 7.5, 50 mM KCL, 0.02 mM EDTA and EGTA, 0.1% BSA and 5 mM Succinate, Sigma) was added to each well with a final digitonin (Sigma #D5628) concentration of 0.002%. BH3 peptides were added at appropriate concentrations and incubated for 1h at room temperature. The cells were then fixed with 15 µl of 8% paraformaldehyde for 15 mins at room temperature. The wells were aspirated to 20 µl using the BioTek 406EL plate washer (BioTek) and neutralized with 20 µl of Tris/Glycine buffer (1.7 M Tris, 1.25 Glycine pH 9.1, Sigma) for 5 mins at room temperature. The wells were aspirated to 20 µl using the BioTek 406EL plate washer (BioTek). Cells were stained with 10 µl of antibody staining solution (per well: 0.0075 µl of cytochrome c-Alexa647 antibody (#612310; 1:2000; Biolegend, to calculate the % of cytochrome *c* positive cells), 0.015 µl of Hoechst 33342 (1:2000; Life Technologies, stains nuclei to calculate the total number of cells), 3 µl of 10X staining solution (10% BSA, 2% Tween20 in PBS, Sigma) and 7 ul of PBS) overnight at 4°C. The next day the cells were washed with BioTek plate washer and imaged. All imaging was performed on the ImageXpress Micro Confocal High-Content Microscope (Molecular Devices; at the Laboratory for Functional Precision Medicine (LFPM) at Harvard Medical School). A 10× objective was used and multi wavelength cell scoring was performed to analyze images using MetaXpress (Molecular Devices; at the LFPM at Harvard Medical School). Cells are scored as negative or positive based on the area. All subsequent analysis was carried out in Excel or GraphPad Prism to calculate cytochrome *c* loss by cells and delta priming scores described in detail in the statistical analysis.

### Flow cytometry-based BH3 profiling

Cells were washed with PBS and stained with zombie-NIR dye (biolegend) for 20 mins at room temperature and washed with PBS. Cells were subsequently stained with anti-human PD-L1-FITC or anti-human EGFR-PE/Cy7 antibodies (Biolegend) for additional 10 mins at room temperature and washed with PBS. Cells were seeded at a density of 1 x 10^4^ in 10 ul of MEB buffer (150 mM mannitol, 10 mM HEPES-KOH pH 7.5, 150 mM KCl, 1 mM EGTA, 1 mM EDTA, 0.1% BSA [bovine serum albumin], and 5 mM succinate) in a 384-well plate (Corning, cat#3575). 10 ul of BIM peptide was added to each well at increasing concentrations (0.001 uM-100 uM) containing digitonin in MEB buffer (0.002%) for 1h at 25°C. Cells were fixed with 5 ul of 10% formalin for 15 minutes followed by neutralization with 5 ul N_2_ buffer (1.7M tris, 1.25M glycine, pH 9.1) for 5 mins at room temperature. 10 ul of cytochrome *c* antibody staining solution was added to each well and incubated overnight at 4°C (antibody staining solution: 1X BD perm/wash solution (BD Biosciences cat#51-209-1KZ) and AF-647-cytochrome *c* antibody (BioLegend cat#612310) diluted 1 in 1000 with an overall final dilution of 1:4000 in the well). Cells are acquired on the iQue flow cytometer the next day and data was analyzed on the iQue software to determine the number of cells that lost cytochrome *c* following 1h exposure to BIM peptide which is indicative of sensitivity to mitochondrial apoptosis.

### CellTox Assay

Parental tumor cells were seeded and at a density of 1 x 10^3^ cells in 100 µl and persister cells were seeded at a density of 5 x 10^3^ cells in 100 µl media in a flat bottom 96 well plate (Corning, catalogue #3904) and left to adhere overnight at 37°C, 5% CO_2_. The next day cells were treated with 0.1% DMSO vehicle control or drugged using the D300e digital drug dispenser (Hewlett Packard) and cultured for an additional 72h at 37°C, 5% CO_2_. On the final day digitonin was added to control wells at a concentration of 0.05% for 15 mins to permeabilize all the cells in the well to allow subsequent quantification of the total number of cells per well after a treatment or following no treatment. 50 ul of 1X celltox green dye (Promega, catalogue #G8743) was added to every well including permeabilized control wells for 15 mins at room temperature. Fluorescence was measured using the Tecan Safire^2^ plate reader from Life Sciences at an excitation/emission spectrum of 485/530 nm.

### CellTiter-Glo® Luminescent Cell Viability Assay

Tumor cells were seeded and a density of 1 x 10^3^ cells in a 96 well plate (Coastar, catalogue #3610) and left to adhere overnight at 37°C, 5% CO_2_. The next day cells were treated with 0.1% DMSO or drugged using the D300e digital drug dispenser (Hewlett Packard) and cultured for 72h at 37°C, 5% CO_2_. On the final 35 µl of CellTiter-Glo® reagent was added to each well (Promega, catalogue #G7570), incubated at room temperature for 20 mins and the luminescence was measured using the Tecan Safire^2^ plate reader from Life Sciences

### T cell cytotoxicity Assay

Cells were trypsinised, washed with PBS and counted using trypan blue and cell counting slides on the Countess FL3 (Invitrogen, Thermo Fisher Scientific). Parental cells, IPCs and DTPs were resuspended in PBS at a density of 1 x 10^6^/ml, parental cells were then stained with VF405 (Biotium, catalogue #30068) (1 uM) and IPCs and DTPs were stained with VF488 (Biotium, catalogue # 30086) (1 uM) for 15 mins at 37°C. The dye was hydrolyzed by adding an equal volume of media and then centrifuged at 500 x g at room temperature. A mixed cell suspension was prepared containing 0.01 x 10^6^/ml VF405-stained parental cells and 0.07 x 10^6^/ml VF488-stained DTPs or 0.07 x 10^6^/ml VF488-stained IPCs and 1ml of mixed cell suspension was seeded into a 24 well plate and left adhere overnight at 37°C, 5% CO_2_. The next day T cells were added at the appropriate effector to target ratio (E:T) ratio (refer to figure legend) for 24h or 48h. The following day the cells were trypsinsed, washed with PBS, stained with zombie-NIR viability dye (BioLegend) for 10 mins at room temperature followed by sequential staining with CD3-APC (BioLegend) for an additional 10 mins. Cells were washed and fixed with 5% formalin and stored at 4°C before acquisition. Prior to sample acquisition on the BD LSR Fortessa Diva software v 10, 15 ul of AccuCheck Counting Beads (Life Technologies, PCB100) were added to each sample. 1000 beads were acquired for each sample and the number of viable CD3-BV421-negative VF405-stained parental cells and viable CD3-negative VF488-stained IPCs or DTPs was calculated using Flojo software version 10.

### Extracellular Flow cytometry staining

Cells were washed with PBS and stained with zombie viability dye for 20 mins at room temperature. Cells were washed and blocking solution was added for 30 mins at room temperature (normal rat serum (1 in 20 dilution), rabbit serum and mouse serum (1 in 50 dilution). Cells were stained with anti-mouse PD-L1-FITC, anti-human PD-L1-BV421, anti-human EGFR-PE/Cy7 or isotype control antibodies (BioLegend) for 10 mins at room temperature. Cells were washed, fixed with 1% paraformaldehyde for 10 mins at room temperature, washed again and stored at 4°C until acquisition on the BD LSR Fortessa using Diva software v10. Data was analyzed using Flojo v10.

### Intracellular Flow cytometry staining

Cells were washed with PBS and stained with zombie viability dye for 20 mins at room temperature. Cells were washed, fixed with 10% formalin for 15 mins at room temperature and washed with PBS. Cells were permeabilized and blocked with 100 ul of 1X BD fix/perm solution (BD Biosciences catalogue # 554723) containing normal rat serum (1 in 20 dilution), normal rabbit serum and mouse serum (1 in 50 dilution), human serum albumin and fetal bovine serum (5%) for 30 mins at room temperature. Fluorochrome conjugated antibodies for Mcl-1, Bcl-2 and Bcl-xl (Mcl-1-AF488, #58326S (1 in 400 dilution), Bcl-xl-PE/Cy7, #81965S (1 in 3000 dilution) and Bcl-2-AF647 #82655S (1 in 400 dilution), Cell Signaling) or isotype control fluorochrome conjugated antibodies (anti-rabbit IgG isotype control-AF488 #2975S, anti-rabbit IgG PE/Cy7 #974925 and anti-mouse IgG1 isotype control-AF647) were added. Cells were incubated overnight at 4°C. The next day, cells were washed with 1X BD fix/perm solution and resuspended in 250 ul of 1X BD fix/perm solution. Cells were acquired using the BD LSR Fortessa using Diva software v10. Data was analyzed using Flojo v10.

### BrdU ELISA

Cells were seeded at a density of 5 x 10^3^ in 100 ul of media in a flat bottom 96 well plate. A BrdU cell proliferation assay was performed following the manufacturer’s instructions (Cell Signaling Technology #6813). Briefly, BrdU was added to each well at a final concentration of 1X and cells were incubated at 37°C, 5% CO2 overnight. The following day the plates were centrifuged and the supernatant was aspirated. Cells were fixed and denatured using 100ul of fixing/denaturing solution for 30 mins at room temperature. Solution was removed and plates were washed 3 times with 1X washing buffer. Secondary antibody was added at a 1X concentration and plates were incubated for 30 mins at room temperature. Plates were washed 3 times with 1X washing buffer. 100 ul of substrate was added to each well for 15 mins and 100 ul of stop solution was then added. The absorbance was measured at 450 nm using the Tecan Safire2 plate reader from Life Sciences.

### Drug efflux pump activity

EFLUXX-ID Gold multidrug resistance assay kit (Enzo, catalogue #ENZ-51030-K100) was used to assess drug efflux activity of three major drug efflux pumps (MDR1, MRP and BCRP) in DTPs according to the manufacturer’s instructions. Briefly, cells were washed with PBS, resuspended at a density of 1 x 10^6^ cells/ml in PBS. For each sample 250 ul of cell suspension was added to a 5 ml FACs tube, 125 ul of each drug efflux inhibitor was added to its respective tube (MDR1 inhibitor, MRP inhibitor or BCRP inhibitor) or 5% DMSO in PBS to untreated or unstained tubes. Cells were incubated for 5 mins at 37°C, 5% CO_2_ and 250 ul of Gold-efflux dye was added to every tube except the unstained and cells were incubated for an additional 25 mins 37°C, 5% CO_2_. Cells were then washed with ice cold PBS and centrifuged for 3 mins at 500 x g at room temperature. Cells were resuspended in ice cold PBS containing PI dye at a final dilution of 1 in 100 and acquired on the BD LSR Fortessa using Diva software v10. Data was analyzed using Flojo v10.

### RNA sequencing

STAR counts used to perform DeSeq2 (Differential sequencing analysis). This is performed relative to a group to determine how genes transcriptionally change e.g IPC vs parental determines if genes in IPC group are up or down relative to the parental. This analysis gives Log2FC and adjusted p-value = these are used for volcano plots (all genes) and also significant genes from here are used to plot Venn diagrams. Gene set enrichment analysis (GSEA) performed using hallmarks pathways for comparison groups e.g (5-FU DTP vs parental. This generated normalized enrichment score (NES) which is either + for an activated pathway corresponding to differentially upregulated genes or – for a repressed pathway corresponding to differentially downregulated genes. Over representation analysis (ORA) uses the genes imported into Enrichr which then ranks them against hallmarks pathways. The representation of genes within a pathway generates an adjusted p-value which represents how significantly represented the genes are within that pathway. Log2FC cutoff used for up/down genes: >=2 (up), <=-2 (down) as well as adj. p-value <=0.05. Raw data for RNA-seq experiments have been deposited at the relevant NCBI platforms, under the accession number GSE308647.

### Statistical Analysis

Please refer to the legend of the figures for description of sample sizes and statistical tests performed. To compare between two groups a paired t-test was used, to compare between three or more groups a one-way ANOVA was used. Data were plotted and analyzed with GraphPad Prism version 10 software. Differences were considered statistically significant when the p value was less than 0.05, and otherwise not significant (ns). BIM BH3 peptide dose response area under the curve (AUC) for cytochrome *c* release was calculated using GraphPad Prism version 10. The variance is similar between the groups that are being statistically compared. Sample size was chosen based on the literature to ensure adequate power to detect a pre-specified effect size.

## Results

### Tumor cells that persist following CAR T cell attack acquire a reduced sensitivity to several drug types and radiotherapy

CAR T cells often kill tumor cells via induction of mitochondrial apoptosis, which is also employed by many chemotherapy agents and radiotherapy^14,23,24^. It is plausible, therefore, that tumor cells subjected to CAR T cell attack would select for reduced sensitivity to CAR T cell-induced mitochondrial apoptosis, which might then translate to a reduction in sensitivity to other apoptosis-inducing agents. Therefore, we tested if tumor cells that survive CAR T cell attack might also be less sensitive to standard chemotherapy agents and other more targeted drugs such as an epigenetic inhibitor (panobinostat, pan-HDACi) and a multi-cell cycle inhibitor (dinaciclib, CDK-5/7/9 inhibitor) as well as radiotherapy.

To create IPCs we co-cultured EGFR-CAR T cells with the Hela or U251 cell line for 3 weeks, refreshing the CAR T cells every 3 days (Fig 1A). We first confirmed that the Hela and U251 IPCs were less sensitive to CAR T cell attack by flow cytometry (Fig 1B-D). We also confirmed that a therapy holiday for several weeks allowed the IPCs to regain sensitivity to EGFR-CAR T cell killing consistent with the theory that persister cells regain sensitivity to the therapy they survived if given a therapy holiday^1,5,25–28^ (Supplemental Figure 1B-C). To test the sensitivity of IPCs to pharmacological agents and radiotherapy we utilized a CellTiter-Glo assay first. We observed that Hela IPCs were less sensitive to several different chemotherapies with distinct mechanisms of action which included 5-FU (anti-metabolite), oxaliplatin (DNA intercalator) and etoposide (topoisomerase II inhibitor) (Fig 1E). Furthermore, we also observed that Hela IPCs were less sensitive to targeted agents such as panobinostat (pan-deacetylase inhibitor) and dinaciclib (multi cyclin-dependent kinase inhibitor (CDKi) of CDK-2/5/9). Similar findings were recapitulated with the U251 IPC cell line (Fig 1F). We confirmed these findings using a Celltox assay in the U251 cell line (Fig 1G). We did observe that dinaciclib killed more U251 IPCs compared to the parental cells (Fig 1G) demonstrated using the CellTox assay which measures the fraction of dead cells using a DNA binding dye that passes through permeabilized cell membranes of dead cells but this wasn’t apparent using the CellTiter-Glo assay which measures overall cell viability by assessing ATP production (Fig 1F). Interestingly, we also observed that both Hela and U251 IPCs were less sensitive to radiation (Fig 1H and 1I, respectively). We saw similar results using the CellTox assay in the U251 (Fig 1J) cell line. Moreover, a reduction in proliferation is a common characteristic observed in DTPs, therefore we investigated if IPCs have a slower proliferative rate compared with parental cells and found that they did (Fig 1K-L). We also investigated if the Hela and U251 IPCs became larger in size and found that their forward scatter-area (FSC-A) which is indicative of the size of a cell had increased compared with the parental cells by flow cytometry (Fig 1M and N). In addition, we also observed that the side scatter area (SSC-A) was increased in IPCs which is indicative of the granularity/complexity of a cell (Fig 1M and N).

**Figure 1:**
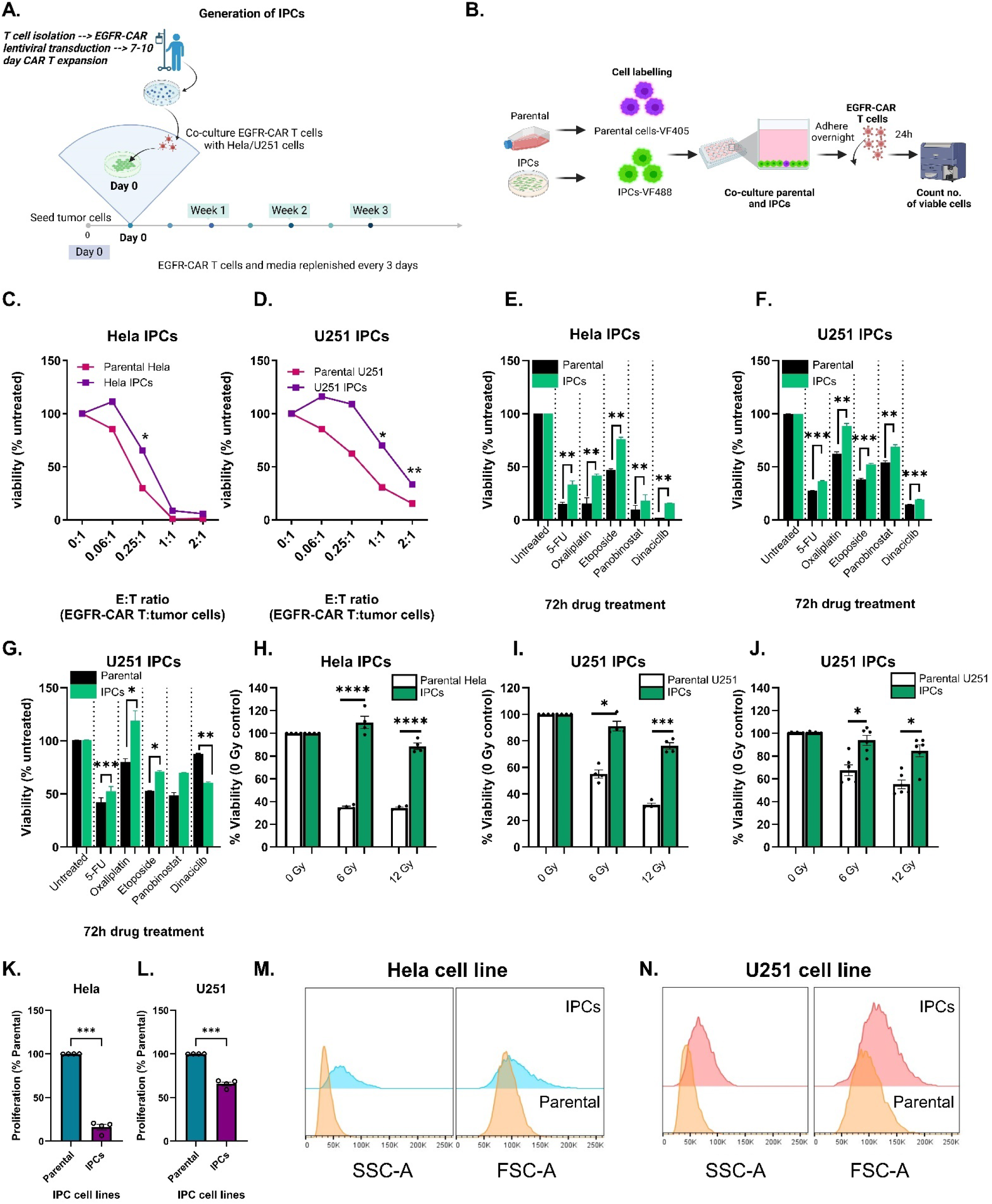
IPCs are less sensitive to chemotherapies and radiotherapy. Schematic showing the generation of IPCs (A) and experimental setup to test sensitivity of IPCs to EGFR-CAR T cell killing (B). Sensitivity of Hela IPCs (C) and U251 IPCs (D) to 24h EGFR-CAR T cell attack, assessed by counting the number of viable cells using counting beads and a dead cell exclusion dye (zombie-NIR) by flow cytometry. A CellTiter-Glo assay was used to assess the viability of Hela IPCs (E) and U251 IPCs (F) following 72h treatment with 5-FU, oxaliplatin, etoposide, panobinostat and dinaciclib. A CellTox assay was also used to assess the viability of U251 IPCs (G) following 72h treatment with 5-FU, oxaliplatin, etoposide, panobinostat and dinaciclib. A CellTiter-Glo assay was utilized to assess the viability of Hela IPCs (H) and U251 IPCs (I) to 6 Gy and 12 Gy irradiation 72h post-irradiation. A CellTox assay was also utilized to assess the sensitivity of U251 IPCs (J) to irradiation. The proliferative rate of Hela IPCs (K) and U251 IPCs (L) was assessed by brdU assay. The SSC-A and FSC-A for parental and IPCs is shown in (M) and (N) for the Hela cell line and U251 cell line model, respectively, determined by flow cytometry. A minimum of n=4 independent biological replicates was conducted. To compare between more than 3 groups a one-way ANOVA, Geisser-Greenhouse correction, Sidak’s multiple comparisons test was used. To compare between 2 groups a paired t-test was used. *p < 0.05; **p < 0.01; ***p < 0.001; ****p < 0.0001.

**Supplemental for Figure 1:**
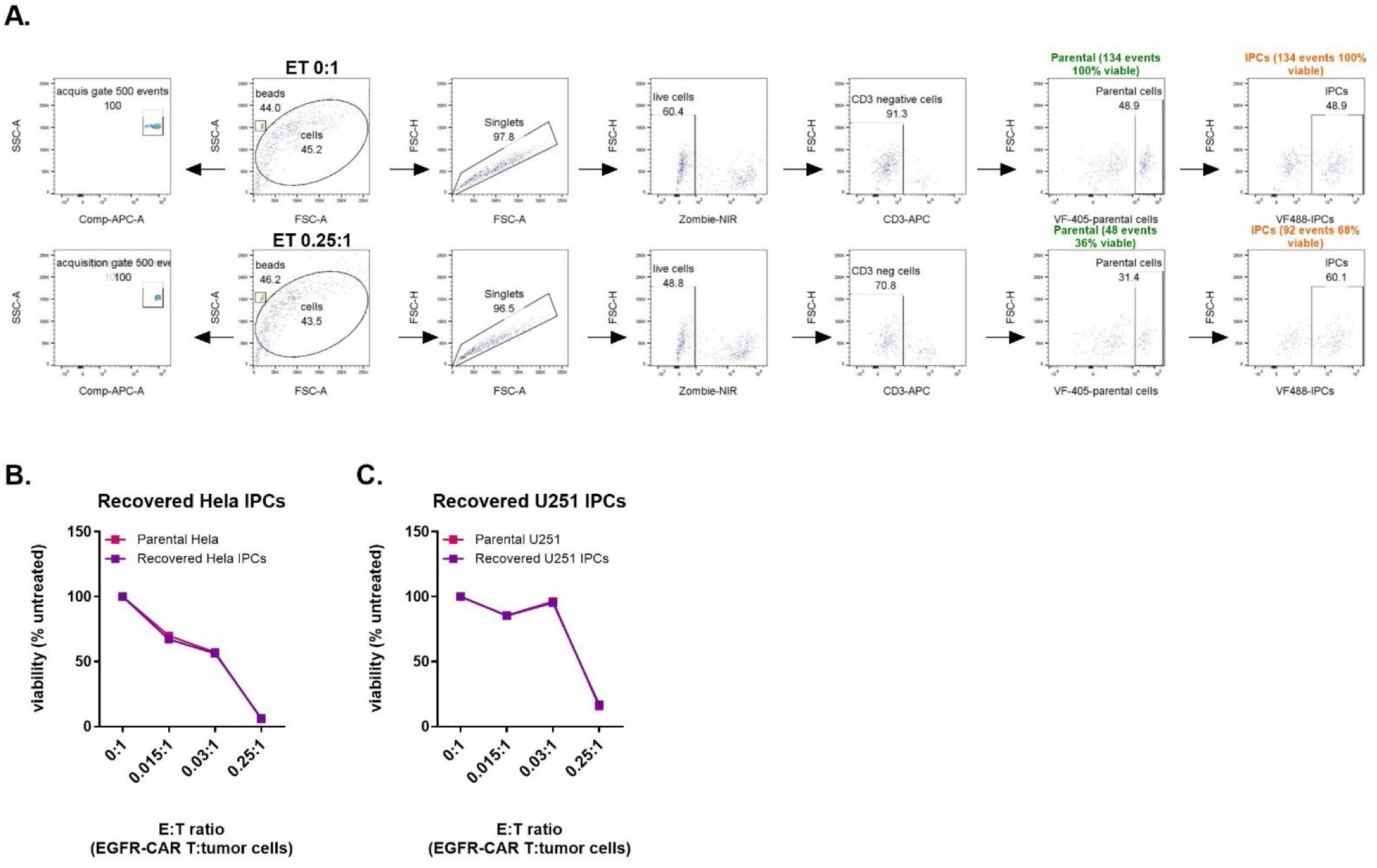
(A) Gating strategy to assess sensitivity of Hela parental cells and Hela IPCs to CAR T cell attack. Parental cells were stained with VF405 and IPCs with VF488 and co-cultured together overnight. EGFR-CAR T cells were added for 24h and viability of VF405-parental Hela and VF488-IPCs were assessed by flow cytometry using counting beads, viability dye (zombie dye) and CD3 antibody. 500 counting beads were acquired for each sample, two samples shown, sample 1 (top row) E:T 0:1 and sample 2 (bottom row) E:T 0.25:1 (bottom row). Viable parental cells were classified using the following gating strategy: single cells → zombie-negative → CD3-negative → VF405-positive. Viable IPCs were classified using the following gating strategy: single cells → zombie-negative → CD3-negative → VF488-positive. (B) Following a therapy holiday IPCs regain their initial therapeutic sensitivity profile. CAR T cells were washed off IPCs and cells were allowed to recover for 3 weeks. Flow cytometry was used to assess sensitivity of recovered Hela IPCs (B) and U251 IPCs (C) to CAR T cell attack. One-way ANOVA, with Geisser-greenhouse correction and Dunnett’s multiple comparisons, Spearman correlative analysis.

In summary, we observed that IPCs which survived CAR T cell attack also acquired a multi-therapy resistance phenotype which afforded them a reduction in sensitivity to several drug classes and radiotherapy.

### Tumor cells that persist following drug therapy acquire a reduced sensitivity to several drug types, radiotherapy and T cell killing

Mechanisms of resistance in DTPs have been documented and include increased expression of DNA damage repair proteins, upregulation of drug efflux pumps and enhancement in fatty acid oxidation^26^. However, it is unknown if the mechanisms of resistance present in DTPs might also confer resistance to other therapies that are not pharmacological agents such as radiotherapy or immune-mediated killing by T cells. Therefore, in this study we examined if DTPs derived from different chemotherapy agents acquired resistance to radiotherapy and CAR T cell killing.

We generated DTPs by continuously treating tumor cell lines for 7 days with an IC_90_ dose of drug (Fig 2A). We selected 3 different chemotherapies with diverse mechanisms of action that include 5-FU, oxaliplatin and etoposide. Other groups have demonstrated that DTPs have a reduced proliferative rate.^25,29^ Therefore, we tested if the DTPs in our study also have a reduced proliferative rate and we found that they did (Fig 2B-D).

**Figure 2:**
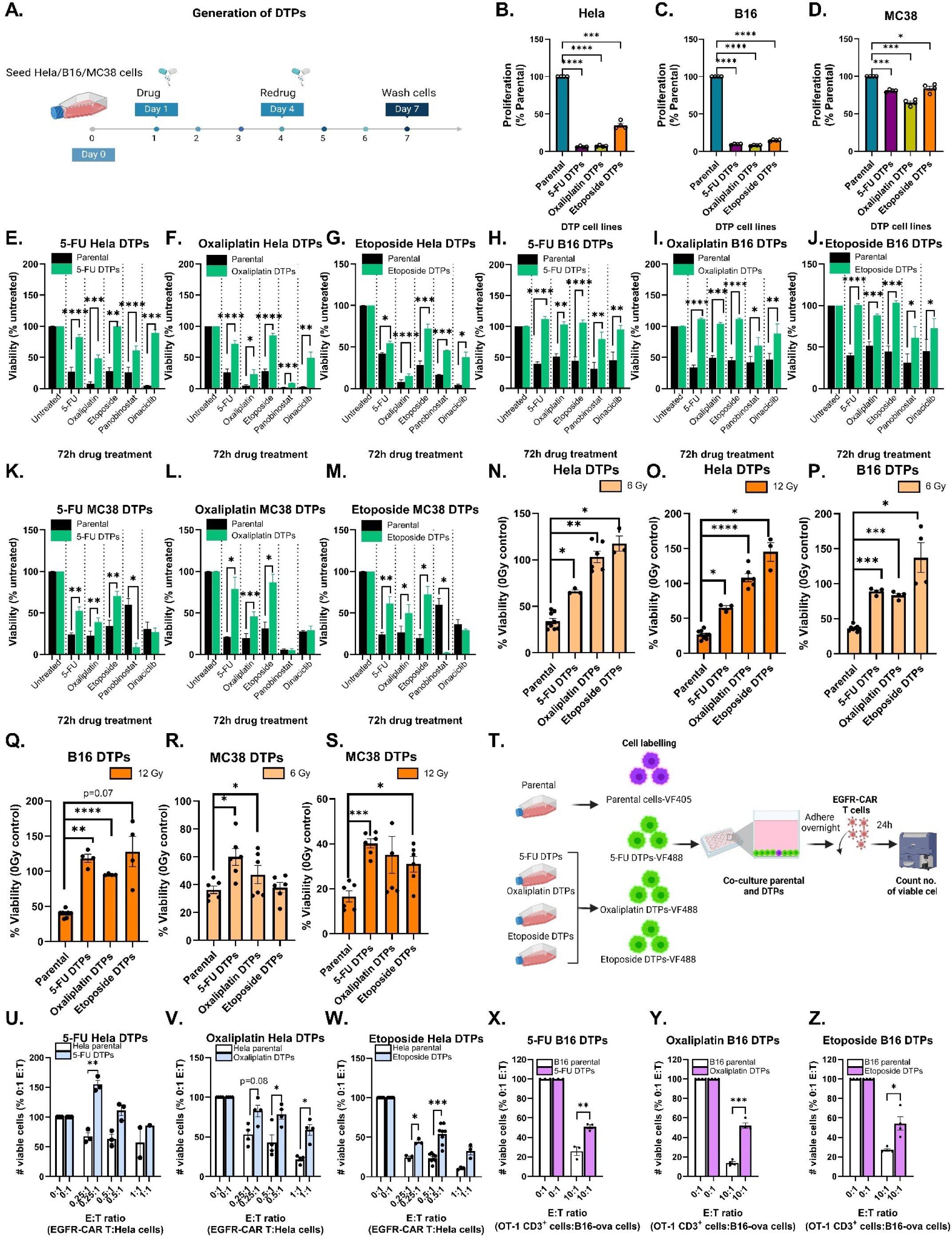
DTPs are less sensitive to chemotherapies, radiotherapy and T cell attack. Schematic showing the generation of DTPs (A). The proliferative rate of Hela, B16 and MC38 DTPs is shown in B-D and was assessed by brdU assay. Sensitivity of 5-FU, oxaliplatin and etoposide DTPs derived from the Hela, B16 and MC38 cell lines to a panel of drugs that included 5-FU, oxaliplatin, etoposide, panobinostat and dinaciclib is shown in E-G (Hela), H-J (B16) and K-M (MC38) respectively, viability was determined by CellTiter-Glo assay following 72h drug treatment. Viability of Hela, B16 and MC38 DTPs was assessed by CellTiter-Glo assay following 72h post 6 Gy and 12 Gy irradiation, data is depicted in N-O (Hela), P-Q (B16) and R-S (MC38). Experimental schematic testing the sensitivity of DTPs to T cell attack is shown in T. Graphical data showcasing the sensitivity of 5-FU, oxaliplatin and etoposide Hela DTPs to EGFR-CAR T cell killing is shown in (U-W) and was determined by counting the number of viable cells using counting beads and a dead cell exclusion dye (zombie-NIR) by flow cytometry. Sensitivity of 5-FU, oxaliplatin and etoposide B16-OVA DTPs to 48h OT-1 CD3^+^ T cell attack is shown in (X-Z). A minimum of n=4 independent biological replicates was conducted, to compare between more than 3 groups a one-way ANOVA, Geisser-Greenhouse correction, Sidak’s multiple comparisons test was used. To compare between 2 groups a paired t-test was used. *p < 0.05; **p < 0.01; ***p < 0.001; ****p < 0.0001.

**Supplemental Figure 2 (First Supplemental for Figure 2):**
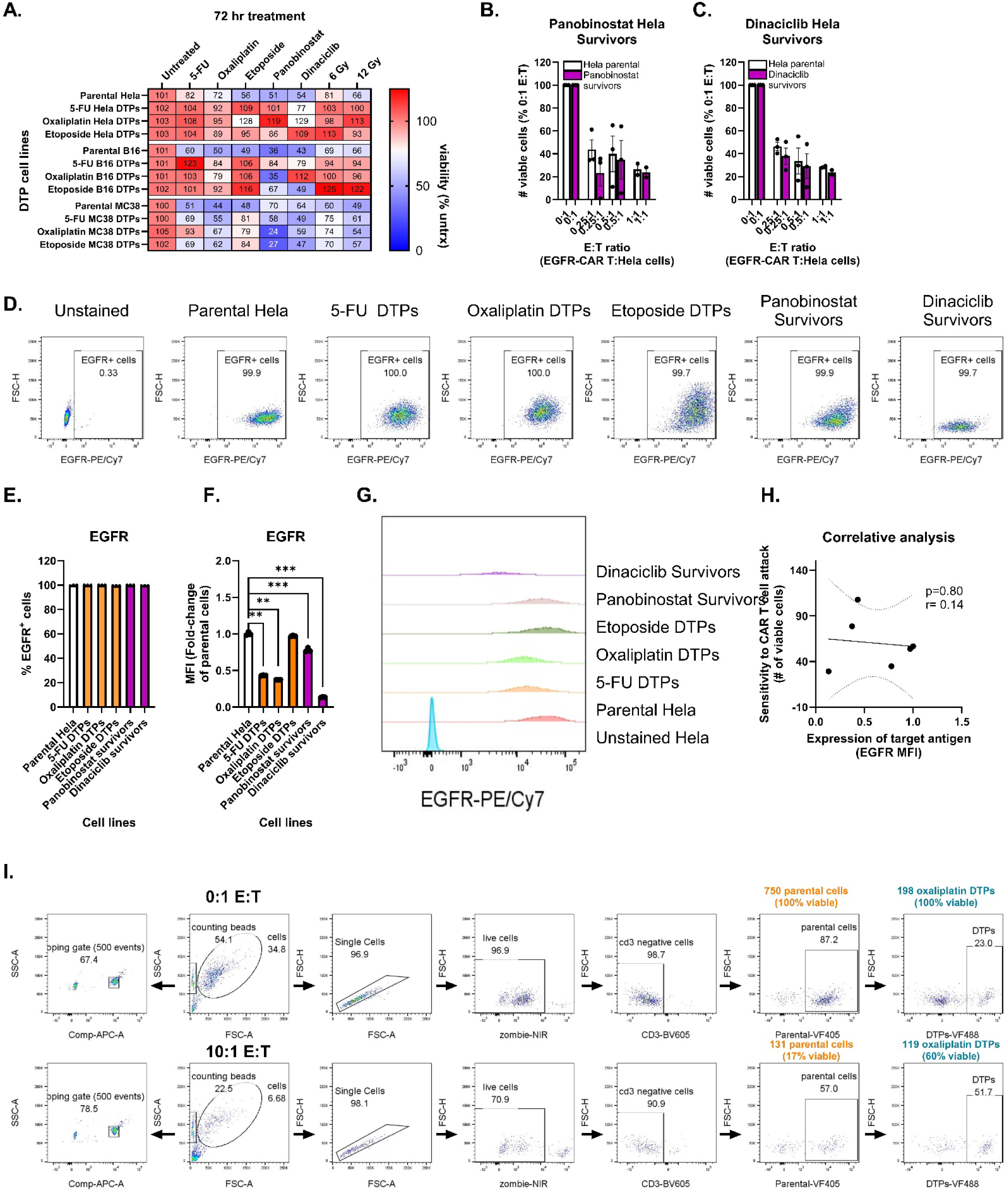
Sensitivity of DTPs derived from the Hela, B16 and MC38 cell lines to a panel of drugs, 6 Gy and 12 Gy irradiation, viability determined using a CellTox assay (A). Sensitivity of Hela cells that survived 7-day treatment with panobinostat (B) and dinaciclib (C) to EGFR-CAR T cell attack, number of viable cells determined by flow cytometry. Percentage of cells positive for target antigen expression, EGFR, determined by flow cytometry is shown in (D) and (E). MFI levels for EGFR, normalized as a fold-change to parental Hela cells is shown in (F) and (G). A correlative analysis of target antigen expression and sensitivity to CAR T cell attack is shown in (H). Gating strategy to assess sensitivity of B16-OVA DTPs to 48h OT-1 CD3^+^ T cell attack (E:T, 10:1) (I). One-way ANOVA, with Geisser-greenhouse correction and Dunnett’s multiple comparisons, Spearman correlative analysis. **p < 0.01; ***p < 0.001.

**Supplemental Figure 3 (Second Supplemental for Figure 2):**
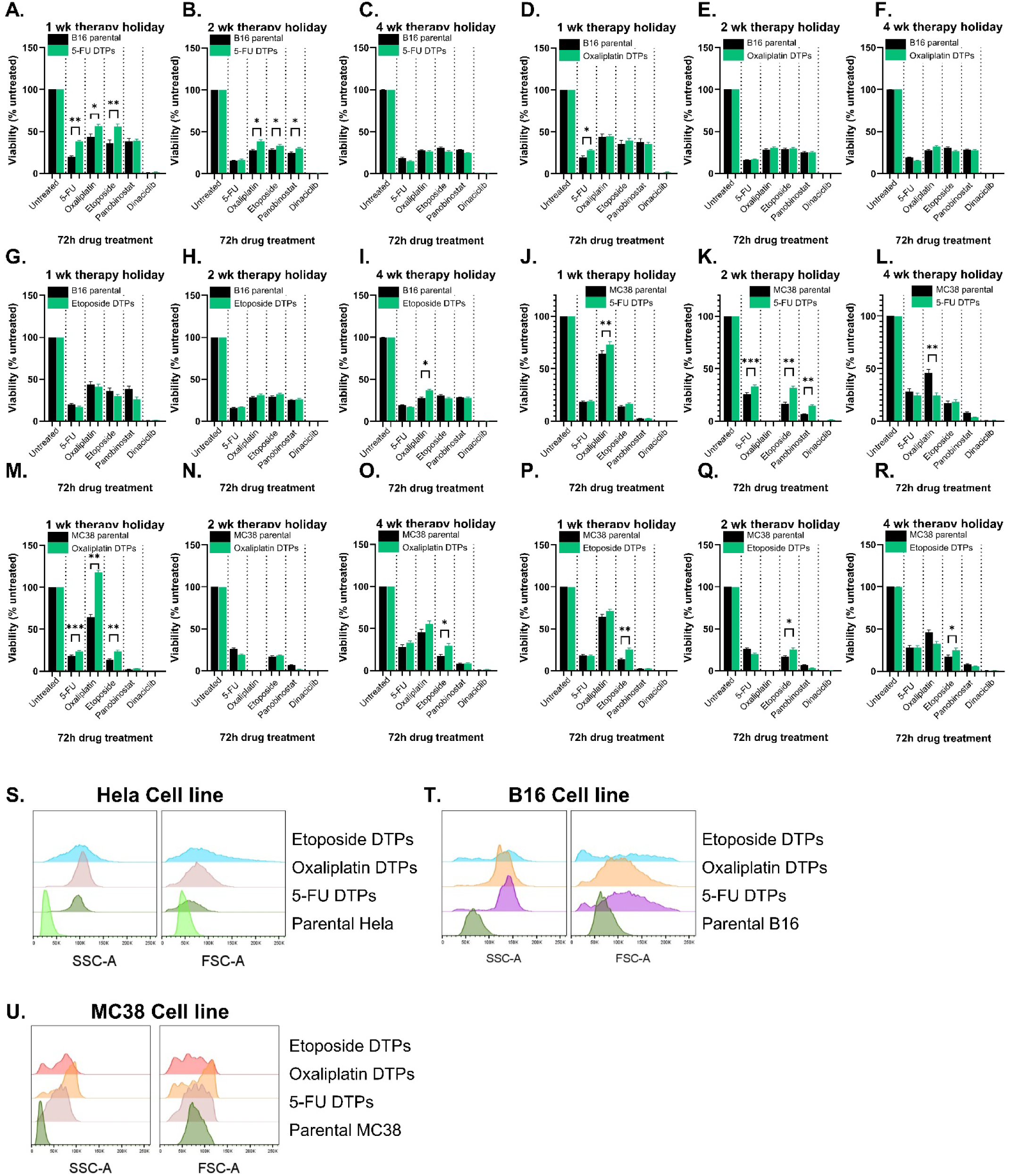
Following a therapy holiday, DTPs regain their initial therapeutic sensitivity profile. Drugs were washed off DTPs and were allowed to recover. CellTiter-Glo assay was used to test sensitivity of recovered 5-FU B16 DTPs (A, D), oxaliplatin B16 DTPs (B,E), etoposide B16 DTPs (C, F), 5-FU MC38 DTPs (G, J), oxaliplatin MC38 DTPs (H, K), etoposide MC38 DTPs (I, L) to pharmacological agents following a 1 week and 4 week drug holiday. SSC-A and FSC-A of DTPs derived from the Hela cell line (M), B16 cell line (N) and MC38 cell line (O) are also depicted, determined by flow cytometry. One-way ANOVA, with Geisser-greenhouse correction and Dunnett’s multiple comparisons, Spearman correlative analysis. *p < 0.05; **p < 0.01; ***p < 0.001.

We observed that 5-FU, oxaliplatin and etoposide Hela DTPs were less sensitive to several chemotherapies that had distinct mechanisms of action (Fig 2E-G, respectively). Similarly, B16 and MC38 DTP cell lines were also less sensitive to these chemotherapies (Fig 2H-J and Fig 2K-M, respectively). However, the sensitivity of the DTPs to targeted therapies such as panobinostat and dinaciclib varied depending on the cell line the DTPs were derived from. For example, DTPs derived from the Hela cell line became more sensitive to panobinostat but remained equally sensitive to dinaciclib (Fig 2E-G). In contrast, 5-FU, oxaliplatin and etoposide B16 DTPs became less sensitive to panobinostat and dinaciclib (Fig 2H-J). DTPs derived from the MC38 cell line became more sensitive to panobinostat and dinaciclib (Fig 2K-M).

A reduction in sensitivity of DTPs to radiotherapy was also observed. Hela and B16 DTPs were less sensitive to 6 and 12 Gy irradiation doses (Fig 2N-O and Fig 2P-Q, respectively). MC38 DTPs developed a small reduction in sensitivity to irradiation (Fig 2R-S). We confirmed these findings using a CellTox assay (Supplemental Figure 2A). We also tested if DTPs were less sensitive to T cell attack (Fig 2T) and found that 5-FU, oxaliplatin and etoposide DTPs derived from the Hela cell line (Fig 2U-W) and derived from the B16-OVA cell line (Fig 2X-Z) were less sensitive to T cell attack.

Similarly to IPCs we found that DTPs also acquired a multi-therapy resistance phenotype and became less sensitive to other drugs, radiation and CAR T cell attack.

### Immunotherapy persister cells acquire a reduction in sensitivity to mitochondrial apoptotic cell death

Considering that IPCs acquire a multi-resistant phenotype and many of these therapeutic agents induce target cell mitochondrial apoptosis, we hypothesized that IPCs may become less sensitive to mitochondrial apoptotic cell death. We assessed IPC sensitivity to mitochondrial apoptosis using microscopy-based BH3 profiling (Fig 3A). We observed that IPCs derived from the Hela and U251 cell line were less sensitive to mitochondrial apoptotic cell death, indicated by a significantly lower percentage of IPCs losing their cytochrome *c* compared with their parental counterparts (Fig B-C, respectively). To gain molecular insight behind why IPCs appear to be less sensitive to mitochondrial apoptotic cell death we hypothesized that one explanation might be an increase in the expression of anti-apoptotic proteins in IPCs. We observed that the levels of Mcl-1, Bcl-2 and Bcl-xl anti-apoptotic proteins were significantly increased in IPCs compared with parental controls in the Hela cell line (Fig 3D-G) and the U251 cell line (Fig 3H-K).

**Figure 3:**
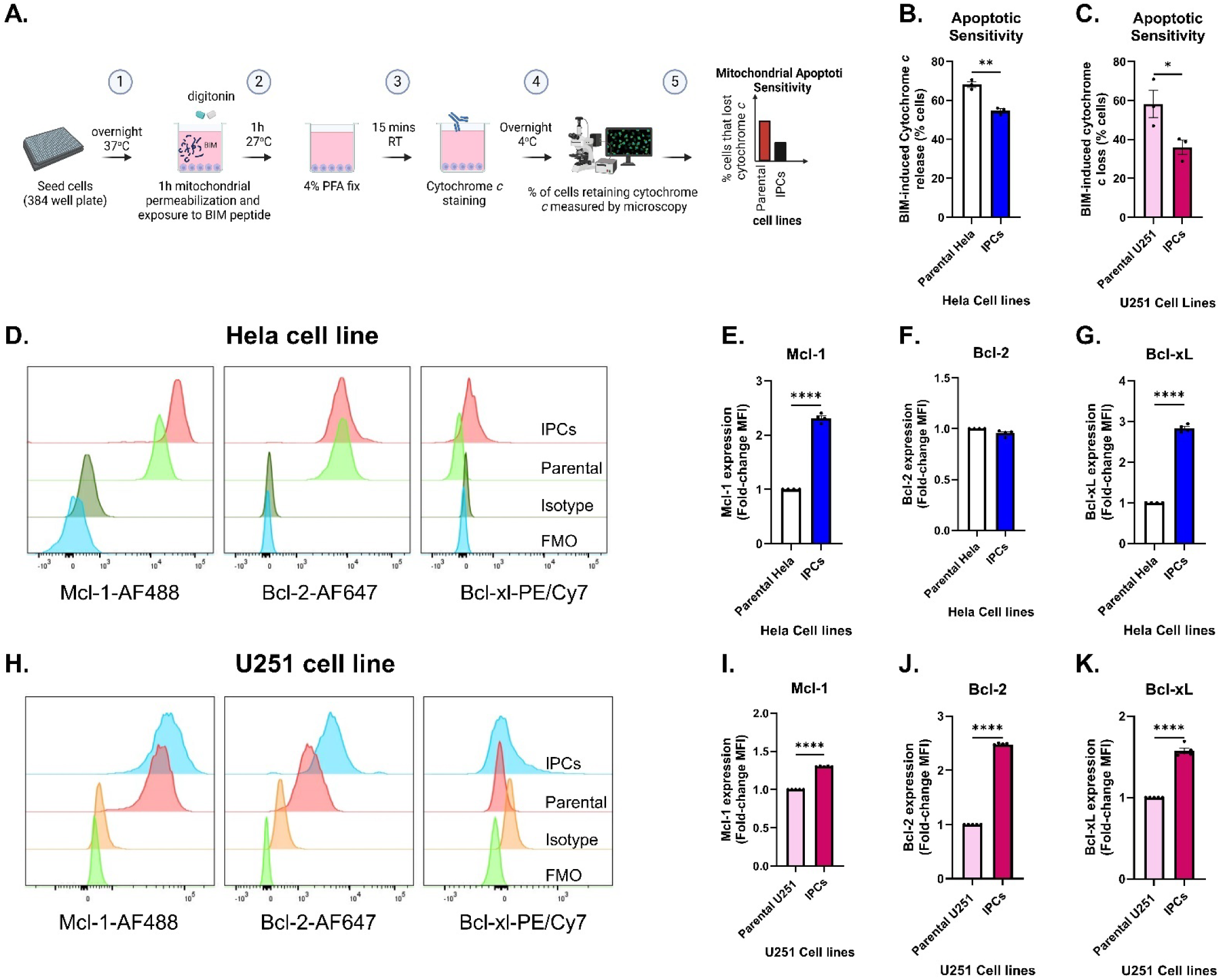
Immunotherapy persister cells acquired a reduction in sensitivity to mitochondrial apoptotic cell death. Experimental schematic of microscopy-based BH3 profiling is shown in (A). BIM-induced cytochrome *c* loss in parental cells and IPCs in the Hela cell line model (B) and U251 cell line model (C). Levels of Mcl-1, Bcl-2 and Bcl-xl anti-apoptotic proteins in parental cells and IPCs, determined by intracellular flow cytometry is shown in (D-G) for the Hela cell line model and (H-K) for the U251 cell line model. Statistical analysis by paired t-test to compare between 2 groups or by one-way ANOVA to compare between two or more groups, with Geisser-greenhouse correction and Dunnett’s multiple comparisons. *p < 0.05; **p < 0.01; ***p < 0.001; ****p < 0.0001.

It is highly plausible that cells rely on multiple mechanisms of resistance to evade CAR T cell attack. A well-documented mechanism of resistance to CAR T cell killing is loss of target antigen. Approximately 50% of the Hela (Fig 4A) and U251 IPCs lost EGFR expression (Fig 4B-C). We theorized that IPCs which did not lose target antigen expression might have to rely more on other mechanisms of resistance for survival such as a reduction in sensitivity to mitochondrial apoptosis. Therefore, we hypothesized that EGFR-positive IPCs would be less sensitive to mitochondrial apoptosis compared to EGFR-negative IPCs. We tested this using flow cytometry-based BH3 profiling (Fig 4D) in the Hela IPC cell line model. We found that EGFR-positive Hela IPCs were less primed for mitochondrial apoptotic cell death compared with EGFR-negative Hela IPCs (Fig 4E). Another well-documented mechanism of resistance to CAR T cell killing is an upregulation of PD-L1 on the surface of tumor cells, we also observed that approximately 50% of Hela IPCs upregulated PD-L1 (Fig 4F-H). Similarly, we postulated that IPCs that are negative for PD-L1 expression would be more vulnerable to CAR T cell attack and be forced to rely on other mechanisms of resistance for survival, therefore we tested if PD-L1-negative IPCs were less sensitive to mitochondrial apoptotic cell death and we found this to be true (Fig 4I).

**Figure 4:**
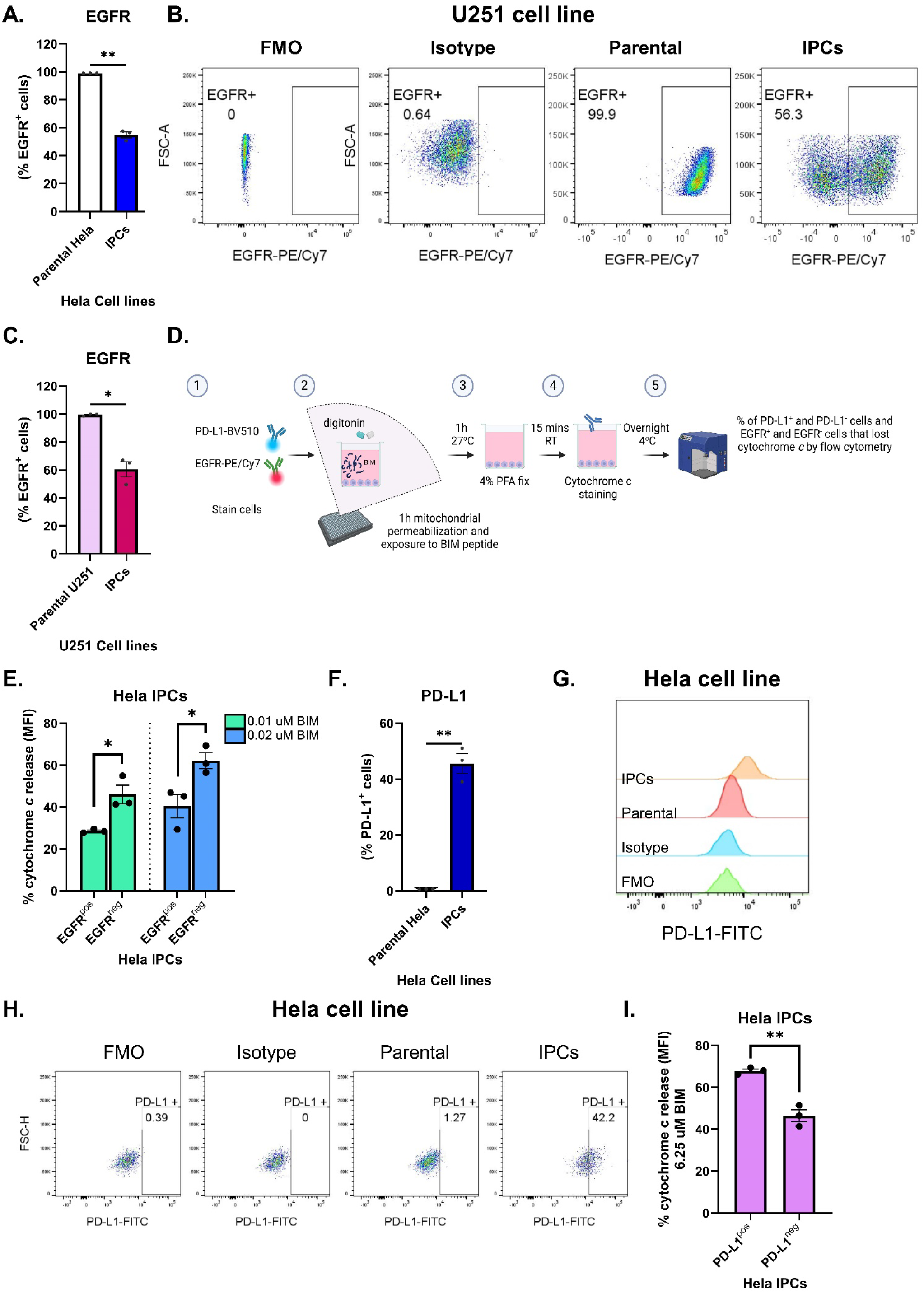
A greater reduction in sensitivity to mitochondrial apoptotic cell death was observed in the immunotherapy persister cells that did not downregulate antigen or upregulate PD-L1. The percentage of parental cells and IPCs positive for EGFR target antigen expression is shown in (A) for the Hela cell line model and (B-C) for the U251 cell line model, determined by flow cytometry. Experimental schematic of flow cytometry-based BH3 profiling is shown in (D). Bim-induced cytochrome *c* loss (MFI) in EGFR^positive^ Hela IPCs and EGFR^negative^ Hela IPCs is shown in (E). Percentage of parental Hela cells and Hela IPCs positive for PD-L1 expression is shown in (F-H). Bim-induced cytochrome *c* loss (MFI) in PD-L1^positive^ Hela IPCs and PD-L1^negative^ Hela IPCs is shown in (I). Statistical analysis by paired t-test to compare between 2 groups or by one-way ANOVA to compare between two or more groups, with Geisser-greenhouse correction and Dunnett’s multiple comparisons. *p < 0.05; **p < 0.01; ***p < 0.001; ****p < 0.0001.

**Supplemental Figure 4 (Supplemental for Figure 3).**
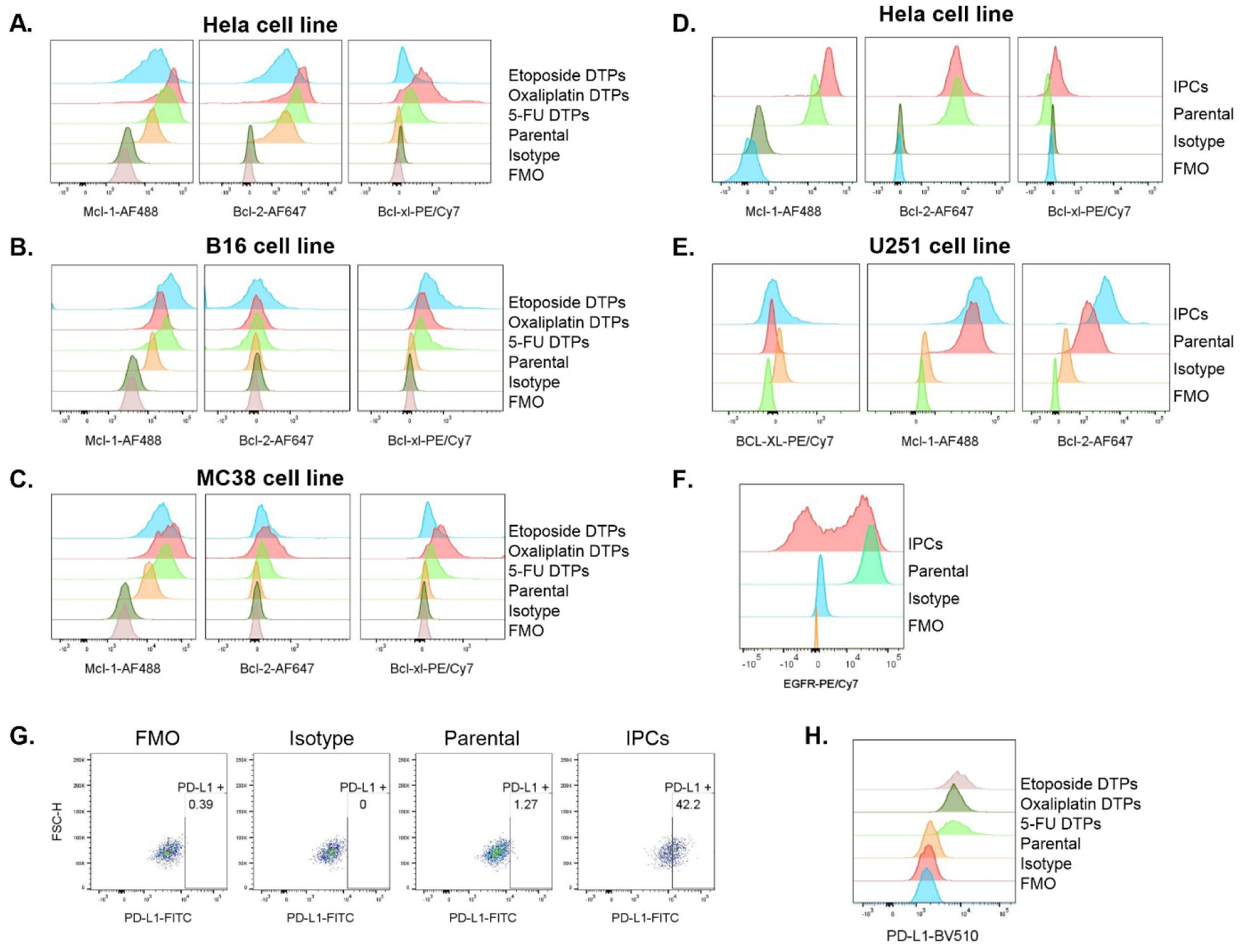
Flow cytometry controls: (A) Representative histograms showing expression of anti-apoptotic proteins in DTP cell lines derived from hela (A), B16 (B) and MC38 (C) cells and in IPCs derived from the hela (D) and U251 (E) cells. Representative histograms showing expression of EGFR (target antigen for EGFR-CAR T cells) in IPCs derived from U251 cells (F). Representative dot plots and histograms showing expression of PD-L1 in IPCs (G) derived from Hela cells and in DTPs derived from B16 cells (H). Fluorescence minus-one (FMO) and isotype controls included for each antibody.

Taken together, we found that IPCs employed several mechanisms of resistance including a decrease in sensitivity to mitochondrial apoptotic cell death, downregulation in target antigen expression and an increase in surface expression of PD-L1. Moreover, the expression of one mechanism of resistance apparently lowered selection pressure for other mechanisms of resistance in the pathway leading to cell death by the mitochondrial pathway of apoptosis.

### Drug tolerant persister cells often acquire an altered apoptotic priming signature

Similarly, we also hypothesized that DTPs might also be less primed for mitochondrial apoptotic cell death as a contributing factor to their multi-therapy resistant phenotype. We observed that Hela, B16 and MC38 DTPs became less primed for mitochondrial apoptotic cell death, indicated by significantly less DTPs losing cytochrome c compared with their parental counterparts in a BH3 profiling assay (Fig 5A-C). To provide some molecular insight behind why DTPs appear to be less sensitive to mitochondrial apoptotic cell death we tested if DTPs have increased levels of anti-apoptotic proteins. This is one potential explanation; other changes could include post-translational modifications which we did not investigate. In the Hela cell line model, we observed that Hela DTPs had elevated levels of Mcl-1, Bcl-2 and Bcl-xl anti-apoptotic proteins compared to the parental cell line (Fig 5D-G). We saw similar results in the B16 and MC38 DTP cell line models (Fig 5H-K and 5L-N).

**Figure 5:**
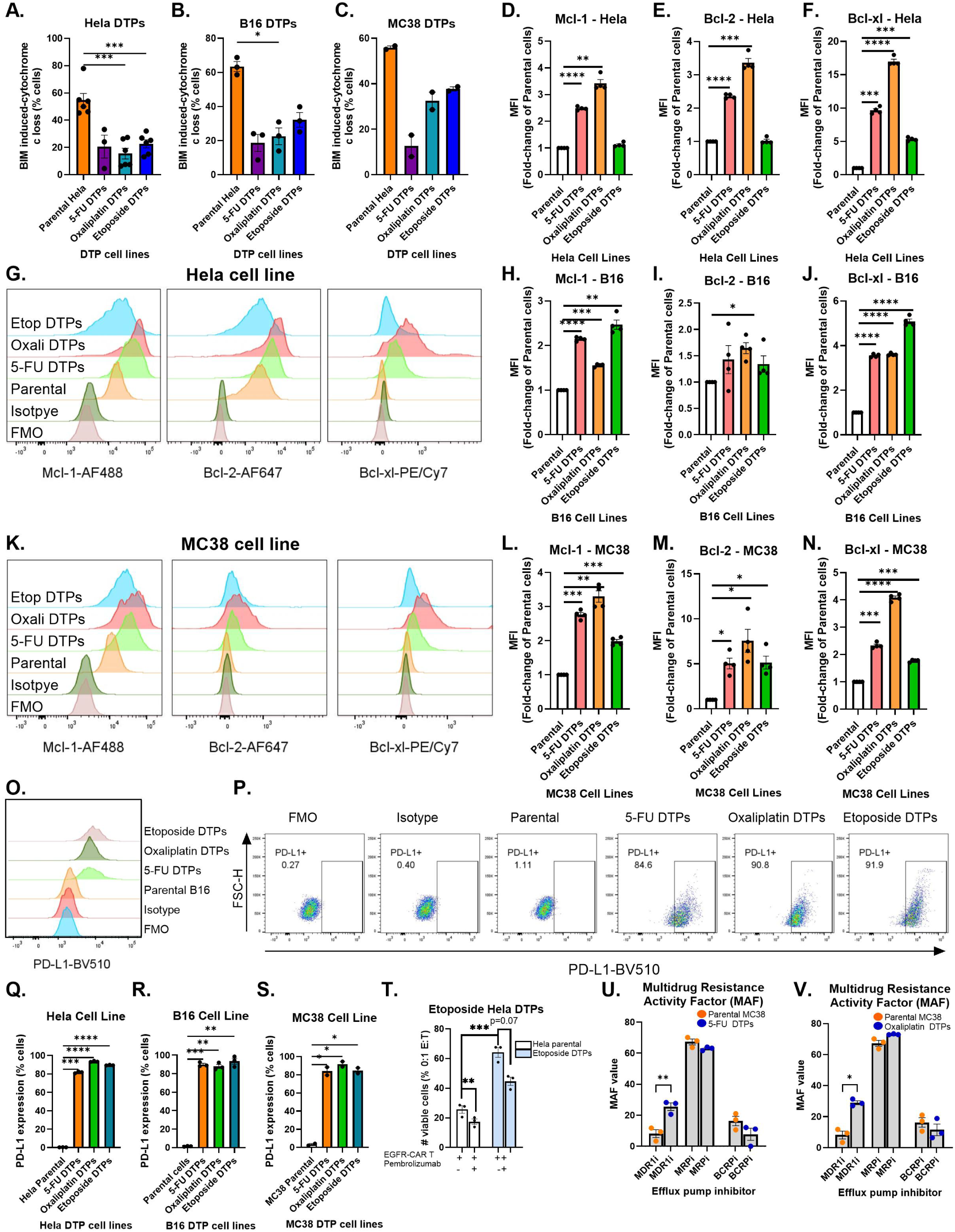
Drug tolerant persister cells acquire an altered apoptotic priming signature. BIM-induced cytochrome *c* loss in parental cells and DTP cells in Hela (A), B16 (B) and MC38 (C) cell line models, measured by microscopy. Levels of Mcl-1, Bcl-2 and Bcl-xl anti-apoptotic proteins in parental cells and DTPs, determined by intracellular flow cytometry is shown in (D-G) for Hela cell line, in (H-J) for B16 cell line and in (K-N) for MC38 cell line models. Frequency of PD-L1 positive parental cells and DTPs is shown in (Q-P and R) for the B16 cell line model, in (Q) for the Hela cell line model and in (S) for the MC38 cell line model, expression determined by flow cytometry. The effect of pembrolizumab on the cytotoxicity of EGFR-CAR T cell mediated killing of parental Hela and etoposide Hela DTPs is shown in (T), viability determined by dead cell exclusion dye and counting beads by flow cytometry. Drug efflux activity assessed in 5-FU and oxaliplatin MC38 DTPs in (U) and (V). Statistical analysis by paired t-test to compare between 2 groups or by one-way ANOVA to compare between two or more groups, with Geisser-greenhouse correction and Dunnett’s multiple comparisons. *p < 0.05; **p < 0.01; ***p < 0.001; ****p < 0.0001.

**Supplementary Figure 5 (Supplemental for Figure 5):**
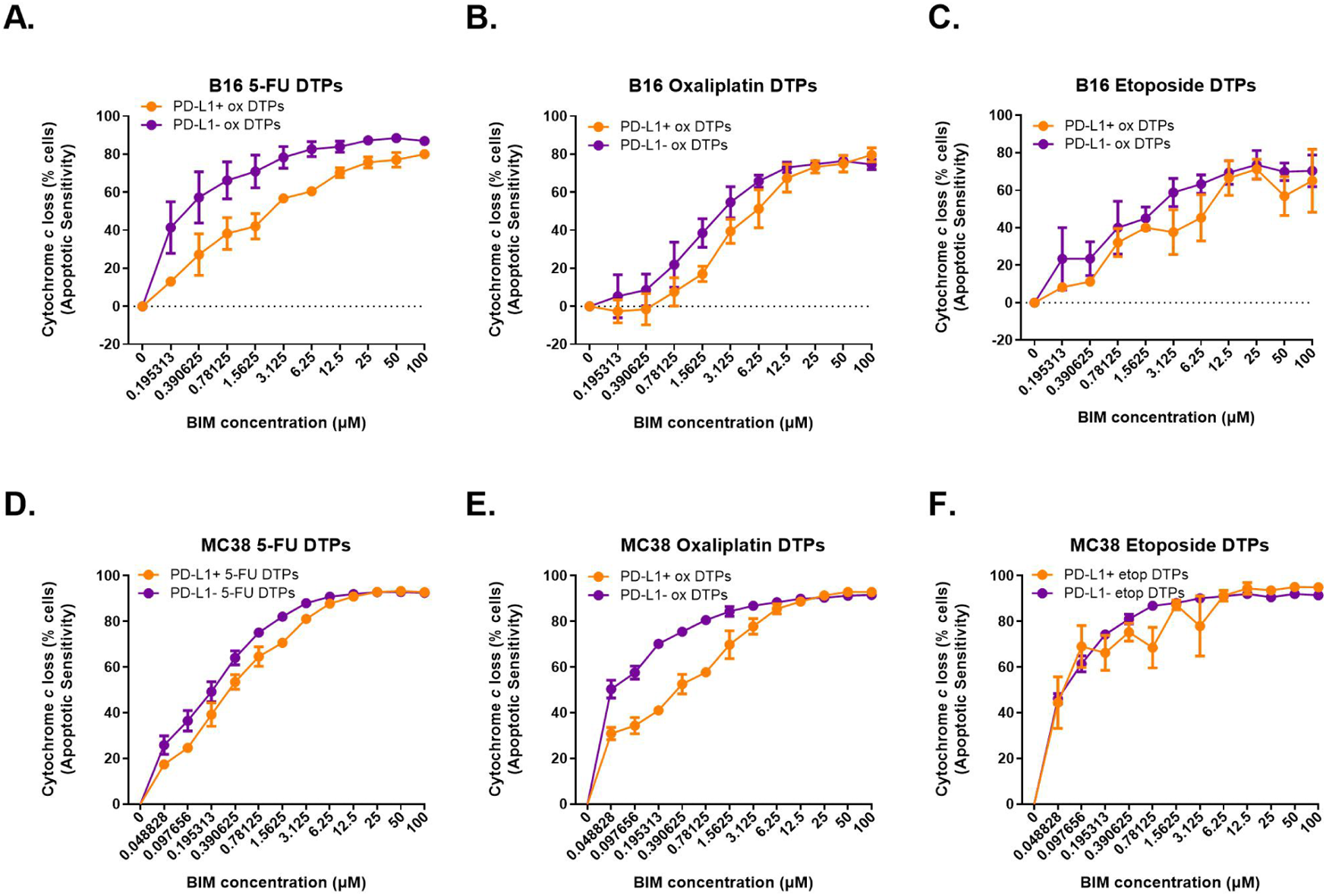
PD-L1-positive DTPs are typically more unprimed for apoptosis compared with PD-L1-negative DTPs. Bim-induced cytochrome *c* loss (MFI) in PD-L1^positive^ DTPs and PD-L1^negative^ DTPs is shown in is shown in (A-C) for the B16 cell line model and (D-F) for the MC38 cell line model measured by flow cytometry. Statistical analysis by paired t-test to compare between 2 groups or by one-way ANOVA to compare between two or more groups, with Geisser-greenhouse correction and Dunnett’s multiple comparisons. *p < 0.05; **p < 0.01; ***p < 0.001; ****p < 0.0001.

Considering that we observed that IPCs upregulated PD-L1 on their surface we sought to investigate if DTPs might share this phenomenon. We observed that approximately 80% of the DTPs upregulated PD-L1 on their surface in the Hela (Fig 5Q), B16 (Fig 5O-P and 5R) and MC38 (Fig 5S) DTP cell line models compared to their parentals which had very low PD-L1 levels of approximately 1-2%. We postulated that PD-L1 may offer protection to DTPs against CAR T cell attack so we tested if PD-1 blockade (pembrolizumab) would enhance EGFR-CAR T cell killing of etoposide Hela DTPs and we found that it did (Fig 5T). Several studies have shown that tumor cells upregulate PD-L1 on their surface in response to chemotherapy treatment and PD-L1 blockade increases drug sensitivity via immune-independent mechanisms. We sought to investigate if PD-L1^+^ DTPs were less sensitive to mitochondrial apoptotic cell death compared with PD-L1^-^ DTPs. We observed very heterogeneous findings; in some cases PD-L1^+^ DTPs were less primed for mitochondrial apoptosis compared with PD-L1^-^ DTPs (Supplemental Figure 5A-E), but in other cases it was comparable (Supplemental Figure 5F).

Other studies have found that DTPs have enhanced drug efflux activity we also tested if this mechanism of resistance is present in the DTP cell line models in our study, we tested this in 5-FU MC38 DTPs and oxaliplatin MC38 DTPs and observed an increase in drug efflux activity in multidrug resistance (MDR, p-glycoprotein) efflux pump but not in MRP or BCRP efflux pumps (Fig 5U and V). An increase in drug efflux activity doesn’t account for a reduction in sensitivity to irradiation or CAR T cell attack. However, this does support the idea that tumor cells upregulate multiple mechanisms of resistance in tandem to protect themselves from cytotoxic therapies and some of these mechanisms can confer resistance to other therapies that the tumor cell hasn’t been exposed to.

Collectively, we identified that DTPs shared similar mechanisms of resistance to IPCs including a reduction in sensitivity to mitochondrial apoptosis and an increase in surface expression of PD-L1. We also observed that DTPs had increased drug efflux activity.

### Targeting mitochondrial apoptotic priming reduces the survival of IPCs

Our group has demonstrated that the use of BH3 mimetics to exploit anti-apoptotic dependencies in tumor cells to increase apoptotic sensitivity is effective in enhancing NK cell-mediated killing of tumor cells^20^. Another study showed that use of venetoclax (BH3 mimetic) increased the intratumoral effector T cells in combination with PD-1 immune checkpoint blockade in pre-clinical murine studies^30^. Considering these recent findings and our results in this study that IPCs are less primed for mitochondrial apoptotic cell death we theorized that use of BH3 mimetics to increase apoptotic sensitivity could increase the sensitivity of IPCs to CAR T cell killing. First we utilized BH3 profiling to identify anti-apoptotic dependencies present in IPCs and their parental counterparts (Fig 5A) by exposing the mitochondria to different peptides that identify dependencies on specific anti-apoptotic proteins (MS1 peptide = Mcl-1, HRK peptide = Bcl-xl, BAD peptide = Bcl-2, Bxl-xl and Bcl-w and MS1 + HRK peptides = Mcl-1 and HRK dependence). We observed that Hela IPCs acquired an increased dependence on Mcl-1 (Fig 6B), Bxl-xl (Fig 6C) and Bcl-2, Bcl-xl and Bcl-w (Fig 6D). The Hela IPCs had a comparable dual dependence on Mcl-1 and Bxl-xl compared to their parental cells (Fig 6E). We observed an increase in dependence on Bcl-xl and Bcl-2 in the U251 IPCs compared to their parental cells (Fig 6G-H) but we didn’t observe significant changes in dependence probing with the other peptides (Fig 6F and I). Another method our group utilized to identify drugs that could increase apoptotic priming of cells is dynamic BH3 profiling by microscopy (Fig 7A). We found that A1331852 and ABT-263 increased the apoptotic priming of Hela IPCs as well as their parental cells (Fig 7B-C). In the U251 cell line, we saw similar results where A1331852 and ABT-263 increased the apoptotic priming of U251 IPCs as well as their parental cells (Fig 6D-E). We next tested if the use of an apoptotic sensitizer could increase the sensitivity of Hela IPCs and U251 IPCs to CAR T cell killing and we found that it did (Fig 6F-H). Furthermore, we also tested if ABT-263 (inhibits anti-apoptotic activity of Bcl-2, Bcl-xl and Bcl-w) could increase the sensitivity of Hela IPCs and their parental cells to treatment with a drug (5-FU) and we found that ABT-263 did increase the sensitivity of Hela IPCs to 5-FU treatment (Fig 6I).

**Figure 6:**
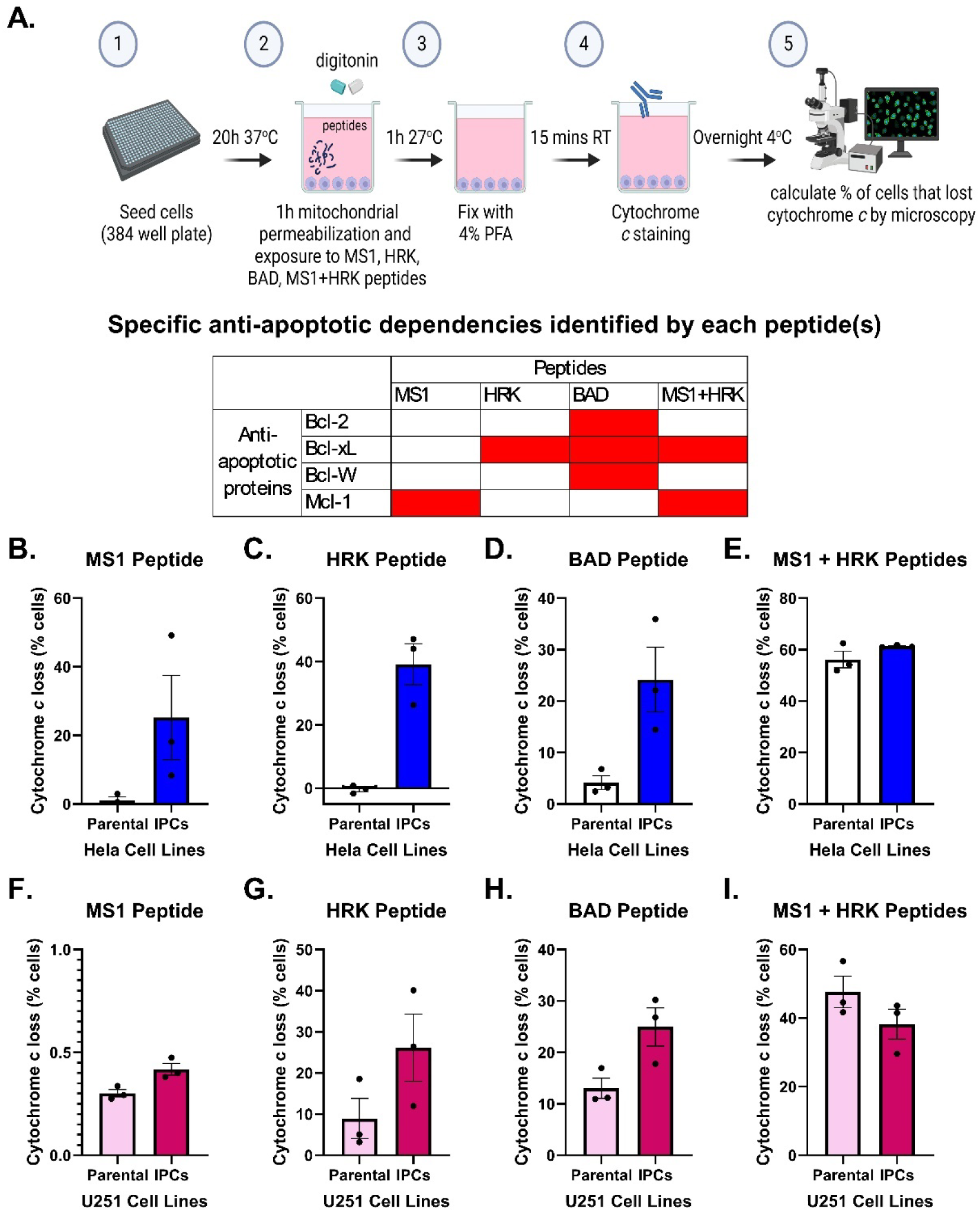
IPCs possess therapeutically targetable anti-apoptotic dependencies that could be harnessed to increase sensitivity to drug and EFGR-CAR T cell attack. Experimental schematic of BH3 profiling assay is shown in (A) which probes for anti-apoptotic dependencies and a table indicating the binding profile of each peptide and their subsequent ability to identify specific anti-apoptotic dependencies. Effect of MS1, HRK, BAD and dual MS1+HRK peptides on cytochrome *c* loss in parental Hela cells and Hela IPCs determined by microscopy-based BH3 profiling (B-E). Effect of MS1, HRK, BAD and dual MS1+HRK peptides on cytochrome c loss in parental U251 cells and U251 IPCs determined by microscopy-based BH3 profiling (F-I). Statistical analysis by paired t-test to compare between 2 groups or by one-way ANOVA to compare between two or more groups. *p < 0.05; **p < 0.01; ***p < 0.001.

**Figure 7:**
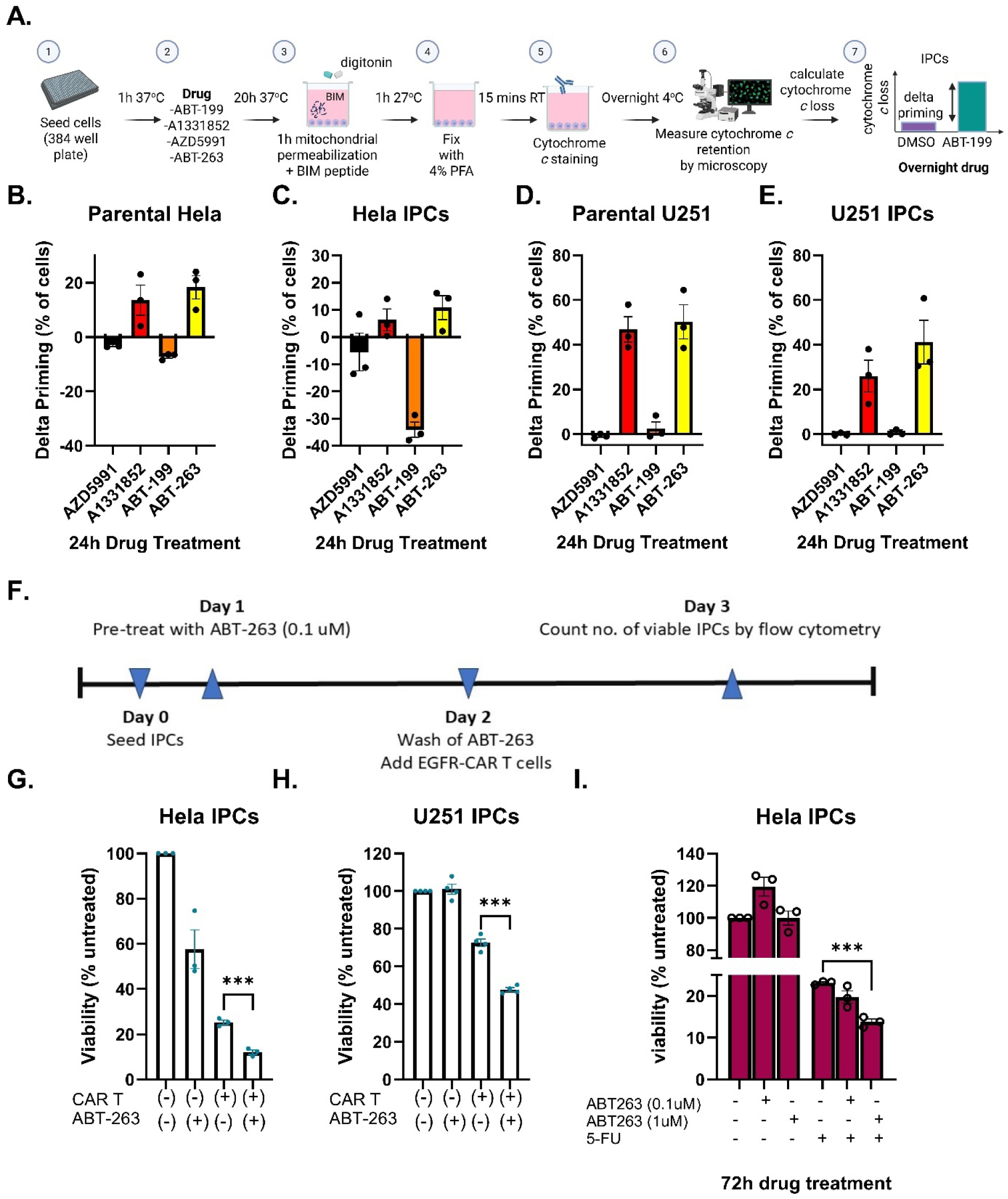
Targeting mitochondrial anti-apoptotic dependencies in IPCs using BH3 mimetics increases their sensitivity to EGFR-CAR T cell attack and drug treatment. Experimental schematic of dynamic BH3 profiling (DBP) assay is shown in (A) which identifies drugs that increases apoptotic priming. DBP data showing the effect of 24h BH3 mimetic treatment on apoptotic priming of Hela parental cells (B) Hela IPCs (C) U251 parental cells (D) and U251 IPCs (E). Experimental schematic for investigating the effect of BH3 mimetic on sensitivity of IPCs to CAR T cell attack is shown in (F) and graphical data is depicted for Hela cell line in (G) and for the U251 cell line in (H), viability determined by dead cell exclusion dye and counting beads by flow cytometry. Effect of ABT-263 BH3 mimetic on sensitivity of IPCs (I) to 5-FU drug treatment, viability assessed by CellTiter-Glo assay. Statistical analysis by paired t-test to compare between 2 groups or by one-way ANOVA to compare between two or more groups. *p < 0.05; **p < 0.01; ***p < 0.001.

Taken together, we observed that targeting anti-apoptotic dependencies in IPCs using BH3 mimetics increased their sensitivity to mitochondrial apoptosis which also translated to an increase in sensitivity to CAR T cell attack and chemotherapy treatment.

### Targeting anti-apoptotic dependencies in DTPs reduced their survival and increased sensitivity to multiple therapeutic modalities

We took a similar approach for the DTPs and hypothesized that increasing the apoptotic priming of DTPs could decrease the survival of DTPs or increase their sensitivity to the drug they survived. First, we employed BH3 profiling to identify anti-apoptotic dependencies in DTPs and their parental cells (Fig 8A). Next we utilized DBP by microscopy to identify BH3 mimetics that could increase the apoptotic priming of DTPs (Fig 8B). In most cases we found that the multi-BH3 mimetic ABT-263 increased apoptotic priming in DTPs in the 3 DTP cell line models. Therefore, we tested if the use of a BH3 mimetic during the 6-day DTP generation process could decrease the fraction of DTPs that survived (Fig 8C). We observed that combining ABT-263 with 5-FU significantly decreased the number of 5-FU B16 DTPs that survived (Fig 8D). We saw similar findings with the 5-FU Hela DTPs (Fig 8E). Next, we asked a different question, would a BH3 mimetic increase the sensitivity of DTPs to the drug they survived (Fig 8F). We observed that ABT-263 increased the sensitivity of 5-FU B16 DTPs to 5-FU, demonstrated by a significant reduction in viability of 5-FU B16 DTPs following a 6-day treatment with 5-FU (Fig 8G).

**Figure 8:**
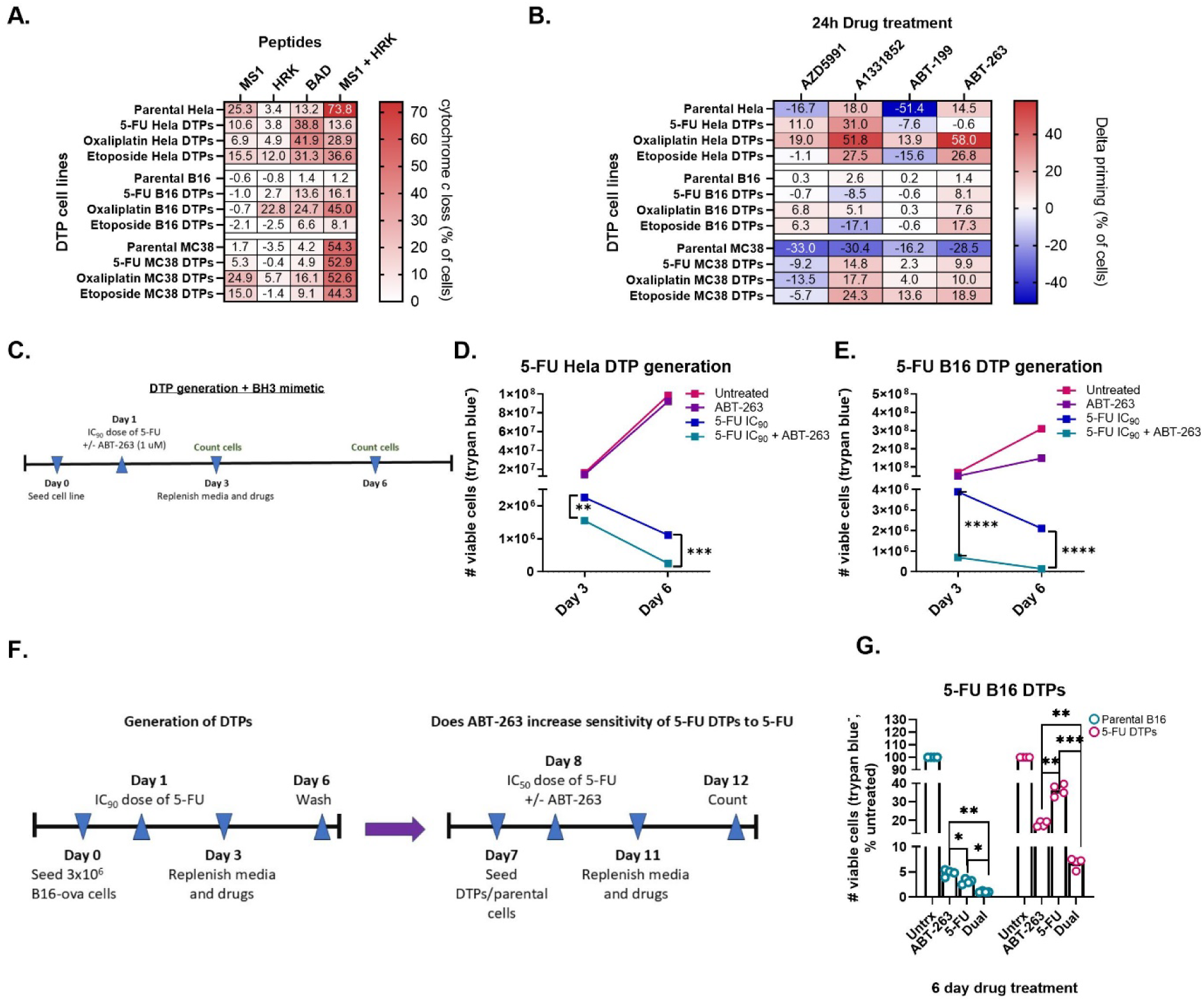
DTPs possess targetable anti-apoptotic dependencies that can be therapeutically harnessed to decrease DTP survival and increase sensitivity to pharmacological agents. Heat map showing effect of MS1, HRK, BAD and dual MS1+HRK peptides on cytochrome *c* loss in parental, 5-FU DTPs, oxaliplatin DTPs and etoposide DTPs in the Hela, B16 and MC38 cell line models shown in (A). Heat map showing the delta priming values (change in apoptotic priming post-BH3 mimetic drug treatment compared with 0 uM drug (DMSO control)) of 4 BH3 mimetics (AZD5991, ABT-199, A1331852, ABT-263) of DTP cell lines is shown in (B). Schematic of experimental set up for D and E is shown in (C). Effect of ABT-263 on the survival of 5-FU B16 DTPs (D) and 5-FU Hela DTPs (E) during the 6-day DTP generation process, number of viable cells quantified using trypan blue and Countess. Experimental set up for G is shown in (F). The ability of ABT-263 to increase the sensitivity of B16 DTPs to 5-FU following a 6-day treatment is shown in (G), viability determined by CellTiter-Glo assay. Statistical analysis by paired t-test to compare between 2 groups or by one-way ANOVA to compare between two or more groups. *p < 0.05; **p < 0.01; ***p < 0.001; ****p < 0.0001.

Similarly to IPCs, targeting anti-apoptotic dependencies in DTPs using BH3 mimetics decreased the survival of DTPs and enhanced their sensitivity to the drug they survived.

### Persisters share similar transcriptional profiles regardless of whether they survived chemotherapy or CAR T cell therapy

Our findings show that regardless of the therapeutic insult, be it an immune cell or a chemotherapy, persisters have similar functional characteristics such as a reduction in sensitivity to programmed cell death and acquisition of a multi-therapy resistant phenotype. This suggests that IPCs and DTPs may not be so different after all. To further test this theory, we carried out bulk RNA sequencing in both IPCs and DTPs using the Hela cell line as a model. The objective of this experiment was to provide an insight into how similar or different the transcriptional changes occurring in IPCs and DTPs are.

We first determined what transcriptional changes were present in the IPCs compared to the parental cells and in the DTPs compared to the parental cells. We identified the genes that are expressed at significantly higher/lower levels and the associated hallmark pathways (GSEA) that are significantly activated/repressed in IPCs (Fig 9A and Fig 9E), 5-FU DTPs (Fig 9B and Fig 9F), oxaliplatin DTPs (Fig 9C and Fig 9G) and etoposide DTPs (Fig 9D and Fig 9H) compared to the Hela parental cells, respectively. The significantly activated hallmark pathways included several different pro-inflammatory pathways and the KRAS signaling pathway. Some of the repressed hallmark pathways included: G2M checkpoint, E2F and Myc pathways.

**Figure 9:**
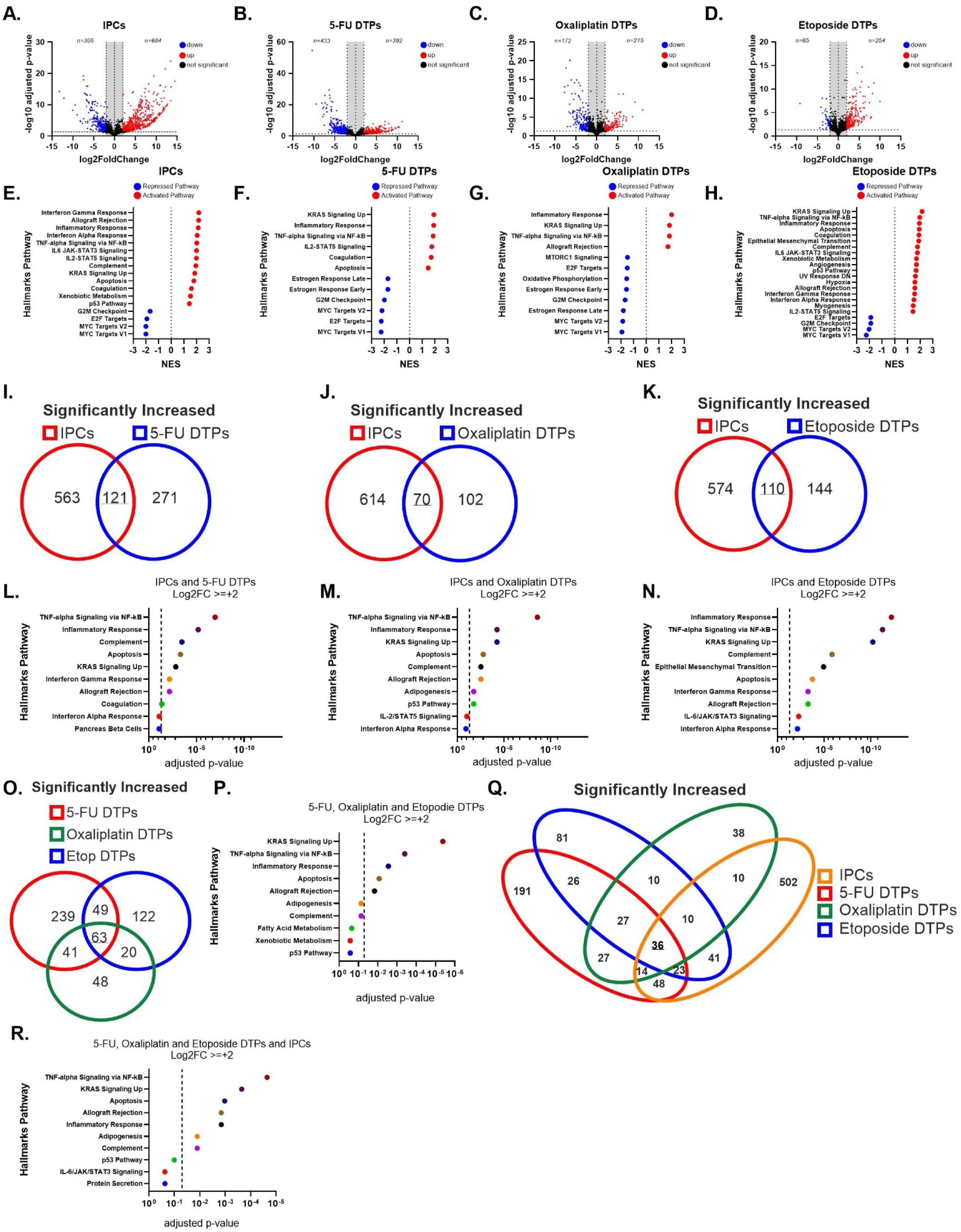
RNA sequencing revealed that pro-inflammatory and KRAS signaling hallmark pathways were upregulated in both IPCs and DTPs. Volcano plots showcasing the differentially expressed genes in IPCs (A), 5-FU DTPs (B), oxaliplatin DTPs (C) and etoposide DTPs (D) compared to the parental Hela cell control. A log2FC cutoff was used as well as an adjusted p-value of <=0.05. Genes that are significantly increased and decreased are shown in red and blue, respectively. A GSEA was conducted on the differentially expressed genes to identify hallmark pathways that are significantly activated (red) or repressed (blue) in IPCs (E), 5-FU DTPs (F), oxaliplatin DTPs (G) and etoposide DTPs (H) compared to Hela parental cells. Venn diagrams outlining the genes that are significantly increased in both IPCs and 5-FU DTPs (I), both IPCs and oxaliplatin DTPs (J) and both IPCs and etoposide DTPs (K). GSEA was conducted on the shared elevated gene expression profiles to identify what hallmark pathways are activated in both IPCs and 5-FU DTPs (L), both IPCs and oxaliplatin DTPs (M) and both IPCs and etoposide DTPs (N). A Venn diagram outlining the genes that are significantly increased in all three DTP groups (O) and the corresponding GSEA for the activated hallmark pathways (P). A Venn diagram showcasing the genes that are significantly increased in both the IPCs and all three DTP groups (Q) and the corresponding GSEA for the activated hallmark pathways (R) is also displayed.

Secondly, we determined if there were any similarities/differences in the transcriptional profiles of IPCs and DTPs. We identified what genes are significantly increased and the associated hallmark pathways that are significantly activated in both IPCs and 5-FU DTPs (Fig 9I and Fig 9L), in both IPCs and oxaliplatin DTPs (Fig 9J and Fig 9M) and in both IPCs and etoposide DTPs (Fig 9K and Fig 9N), respectively. The activated hallmark pathways included different pro-inflammatory pathways (TNF, complement) and the pro-survival KRAS signaling pathway. Although there were many genes that were significantly downregulated in both IPCs and DTPs, there were no shared hallmark pathways that were significantly altered based on the overlapping differentially expressed gene data (data not shown).

**Supplementary Figure 6 (Supplemental for Figure 9):**
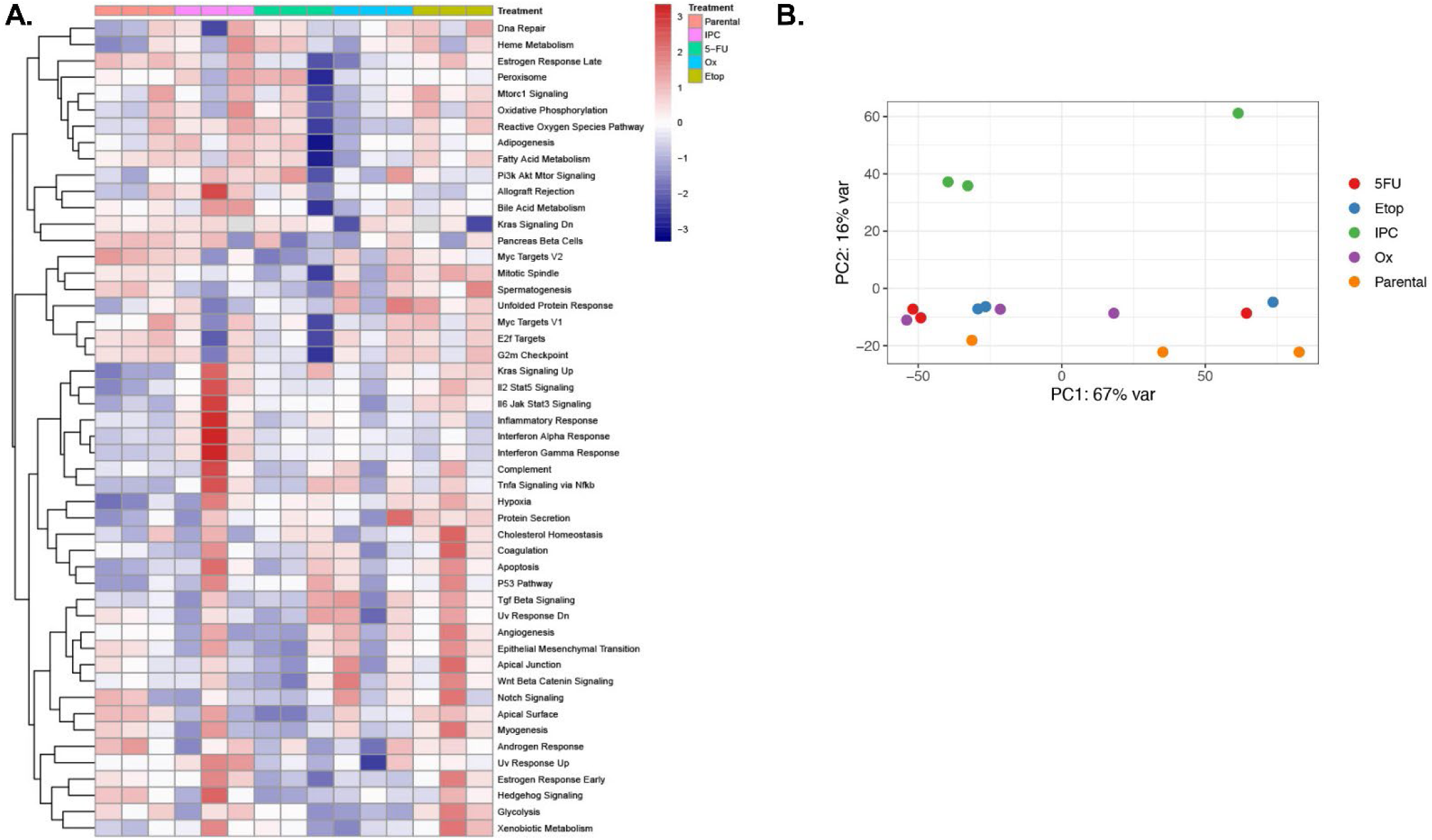
Hierarchical clustering and PCA plots for RNA sequencing data.

Thirdly, we identified the genes that were significantly increased in all three DTPs and the associated hallmark pathways that were significantly activated (Fig O-P) which included: TNF-α signaling via NFκB, KRAS signaling, apoptosis, allograft rejection, inflammatory response. Lastly, we identified 36 genes that were expressed at significantly higher levels in the IPCs and all three DTPs (Fig 9O). GSEA revealed that these gene expression changes resulted in a significant activation in several hallmark pathways (Fig 9R) including TNF-α signaling via NFκB, KRAS signaling, apoptosis, allograft rejection, inflammatory response, adipogenesis and complement.

In conclusion, we observed that there were several hallmark pathways that were significantly activated in all three DTP groups and in the IPCs. These pathways were mainly pro-inflammatory and pro-survival (KRAS signaling).

## Discussion

When a cancer survives following one therapy it often develops insensitivity to other forms of therapies, a phenomenon known as cross-resistance^31^. It remains unclear how cross resistance to therapies that have completely different mechanisms of action occurs. Immunotherapy has become increasingly important in the treatment of cancer in the last decade. Several studies have discovered that tumor cells which survive drug treatment known as drug tolerant persisters (DTPs) also become more resistant to other drug treatments^28^ but it remains unknown if these DTPs become less sensitive to immunotherapy. This prompted us to ask the question, do DTPs also become less sensitive to immune cell killing? Conversely, we also asked the opposite question, could tumor cells that survive immune cell killing (IPCs) become less sensitive to drug treatment? In both cases we found this to be true, DTPs and IPCs developed resistance not only to the therapy they survived but also to other therapies with completely distinct mechanisms of action, recapitulating clinical observations. We began therefore to consider DTPs and IPCs as potentially not completely distinct, but rather as persisters that may share common mechanisms to equip them with multi-therapy resistance.

Immune killing and drug killing of cancer cells are often considered very separately, but our observation of cross-resistance of persister cells suggests that they may not be so different after all. Drugs often kill cancer cells via activating programmed cell death pathways^32^. In the case of the mitochondrial pathway of apoptosis, this can be via the increase in abundance or function of pro-apoptotic proteins relative to anti-apoptotic proteins^32^. While immune cell killing is often simply, and misleadingly, described as “lysis”, immune killing has also been shown to occur via activation of programmed cell death pathways in the target cell^14^. These mechanisms include direct granzyme-mediated cleavage and activation of the pro-apoptotic Bcl-2 family member BID, and direct activation of caspase-3^33,34^, as well as ligation of target cell death ligand receptors in the TNF superfamily^35–37^. From an evolutionary standpoint, this makes sense. If a target cell already possesses rapid and efficient ways of killing itself, why wouldn’t a cytotoxic cell select ways to take advantage of those? Certainly, the mechanisms of initiating apoptotic signaling are very different between drugs and immune cells, but the fact that cells surviving both modalities acquire resistance to apoptosis in surviving cells suggests that from one perspective, they can both be considered as apoptosis induction systems.

The apparent shared use of apoptosis to induce death in cancer cells by both drugs and immune therapy has implications for the rational construction of effective combination therapies. Ni Chonghaile et al^38^, demonstrated that tumors that are less sensitive to mitochondrial apoptosis were less sensitive to drugs, in vitro and in the clinic. Pan et al^15,21,20^ and Pourzia et al^14^ found that insensitivity to apoptotic signaling also conferred resistance to NK cell therapy and CAR T therapy, respectively. In this study, we found that BH3 mimetics that increase mitochondrial apoptotic sensitivity of IPCs and DTPs also increased their sensitivity to CAR T cell attack and drug therapy. Potter et al^39^, Pourzia et al^14^ and Pan et al^20^ have shown that combining pro-apoptotic drugs with another drug or immune cell therapy enhances tumor cell killing, respectively, underlining the significance of sensitivity to mitochondrial apoptotic cell death as a determinant for tumor cell survival in response to a drug or immune cell therapy.

There is appropriate interest in rationally designing effective combinations with immunotherapy in cancer. In most cases, the focus has been on enhancing tumor cell recognition and contact by cytotoxic immune cells^40–43^. Relatively little attention has been paid to the strategy of enhancing the efficiency of tumor cell death once recognition by a cytotoxic cell has been achieved. It is worth noting that killing of a target cell often requires several synapse formation events before the cell commits to apoptosis^20,44,45^. If you consider immune therapies as apoptosis induction systems like drug therapies, designing drug combinations with immune therapies might not be so complicated after all. It might be as simple as combining immunotherapies with any drug that can increase apoptotic sensitivity in tumor cells without killing immune cells. Many clinical trials have tested the combination of immune checkpoint blockers with different types of drug treatments including tyrosine kinase inhibitors, anti-angiogenic therapies, radiotherapy and chemotherapies and in many cases, there is an increase in treatment efficacy^46^. Of course, the immune-stimulatory effects of the drug may contribute to an increase in efficacy^46^. However, it is worth considering that these pharmacological agents might also be increasing the sensitivity of tumor cells to immune cell killing by pushing the tumor cells closer toward the threshold for apoptosis, making them more amenable to immune-mediated killing.

While we have focused here on the importance of insensitivity to apoptotic signaling in cancer cells surviving both drug and immune therapies, it is likely that additional mechanisms that promote cell survival are simultaneously in effect. Some of these mechanisms which been identified in other studies as well as in our study include alterations in drug influx/efflux^47,48^ enhanced DNA repair capacity^49^, upregulation of PD-L1 ^50^, and antigen downregulation^51^. We observed evidence for some of these in this paper along with a reduction in sensitivity to mitochondrial apoptosis. The appearance of decreased apoptotic priming does not rule out the presence of other mechanisms reducing sensitivity. In the case of immune killing, we observed in some IPCs upregulated PD-L1 expression or downregulated antigen expression. It is notable that IPCs that lacked these mechanisms for immune escape were more likely to have a reduced sensitivity to mitochondrial apoptosis. This suggests that the root cause of survival is resistance to apoptosis, which can be effected by changes upstream via upregulation of PD-L1 or downregulation of antigen or proximal to mitochondrial permeabilization downstream.

Interestingly, we also found that DTPs shared another mechanism of resistance with IPCs that included an upregulation of PD-L1 on their surface. Other studies have also shown that chemotherapies upregulate PD-L1^52,53^ and decrease sensitivity to T cell attack^54^, which we also observed to be true in our study in the context of DTPs. Several pro-survival pathways have been implicated in upregulating PD-L1 expression on the surface of tumor cells including MAPK signaling^55^ and EGFR signaling^56^ as well as DNA damage repair signaling (ATM/ATR/Chk1 and STAT1/STAT3)^57^. Therefore, expression of PD-L1 on the surface of tumor cells, in this case DTPs, may identify a population of tumor cells that are more resistant to therapies, with a pro-survival advantage compared to PD-L1-negative tumor cells.

We observed that the IPCs and DTPs have a reduced proliferation rate, repressed G2M and MYC hallmark signaling pathways. These results were consistent with previously published findings showing that persisters have a reduced proliferative rate and suppressed myc signaling^58^. We considered that tumor cells might rely on common pathways to withstand the cellular stress created by the therapy used to generate the persister cell state. We also theorized that this might be independent of the immune or drug therapy used. These theories were supported when we observed that IPCs and DTPs experienced similar transcriptional changes leading to the activation of hallmark pathways characterized by pro-inflammatory signatures and a pro-survival KRAS signaling signature. Results from two studies have indicated that persisters may have an inflammatory profile. Firstly, mitochondrial outer membrane permeabilization (MOMP) is essential for mitochondrial apoptosis, via enabling cytochrome *c* release that leads to caspase activation and rapid programmed cell death^59^. A study by Kalkavan et al, demonstrated that sublethal cytochrome *c* release, facilitated a cell’s entry into the persister cell state following treatment with a BH3 mimetic^60^. Vringer et al, recently demonstrated that MOMP triggers inflammation via NFκB signaling^61^. Collectively, these studies in combination with our work suggest that persister cells may have an inflammatory profile. It has been reported that NFκB promotes chemoresistance via upregulation of anti-apoptotic proteins including Bcl-2 and Bcl-xl which were also upregulated in our persister models^62–64^.

Hela cells homogeneously express the oncogene known as epidermal growth factor receptor (EGFR), which is an upstream activator of KRAS^65^. KRAS signaling promotes cell survival^65^. It has been reported that chemotherapy treatment can reinforce oncogene addiction^65^. Therefore, it might not be that surprising that we observed increased activation of KRAS signaling in both IPCs and DTPs. It is a plausible scenario that a tumor cell might rely even more on an oncogene, becoming further addicted to survive extreme cellular stress while in the persister cell state. This could be a therapeutically exploitable target to eradicate persister cells if a pre-existing oncogene addiction existed in the tumor cells. Further work needs to be carried out to test this hypothesis.

In conclusion, while persistence after immune therapy or after drug therapy are generally treated as very distinct events, we found that persister cells that emerge after these therapies share a common phenotype that is resistant to immune therapy, drug therapy, and radiation. They also share a reduction in mitochondrial apoptotic priming as a mechanism for insensitivity to all these modalities. An important implication is that sensitivity can be restored by enhancing apoptotic priming in both kinds of persister cells. We propose that one principle to enhance the efficacy of immune therapy, as in drug therapy, is to simply identify drugs that enhance apoptotic signaling in target tumor cells that are also tolerable.

## Acknowledgements

This work utilized an Illumina NovaSeq X Plus that was purchased with funding from a National Institutes of Health SIG grant 1S10OD036228-01.

## Conflict of Interest

The authors have no conflict of interest to declare.

## Author Contribution

All authors contributed to this study. M.D. and A.L. designed the study and wrote the manuscript. M.D., C.J.T., D.G., L.B., B.C.C., J.Y-T.K., M.L.O., C.W.W., S.S., L.K., M.Y., G.A., F.B. and A.J.A. performed experiments. J.R., P.B., K.A.S., M.V.M. provided advice on experimental design. D.B., C.P.P., P.H.L., provided cell lines.

## Funding

This work was funded by NCI R01 249062 (AL and MVM). AL also receives funding from Blood Cancer United’s Discovery Grant Program.

## Data Availability

Available upon request.

## Notes

### Competing Interest Statement

The authors have declared no competing interest.

### Summary of Updates

Bulk RNA sequencing of IPCs and DTPs has been conducted to demonstrate that similar transcriptional changes occur in both IPCs and DTPs

