## Supplementary figures and images for "Drug tolerant persisters and immunotherapy persister cells exhibit cross-resistance and share common survival mechanisms"

### Supplemental Figure 1

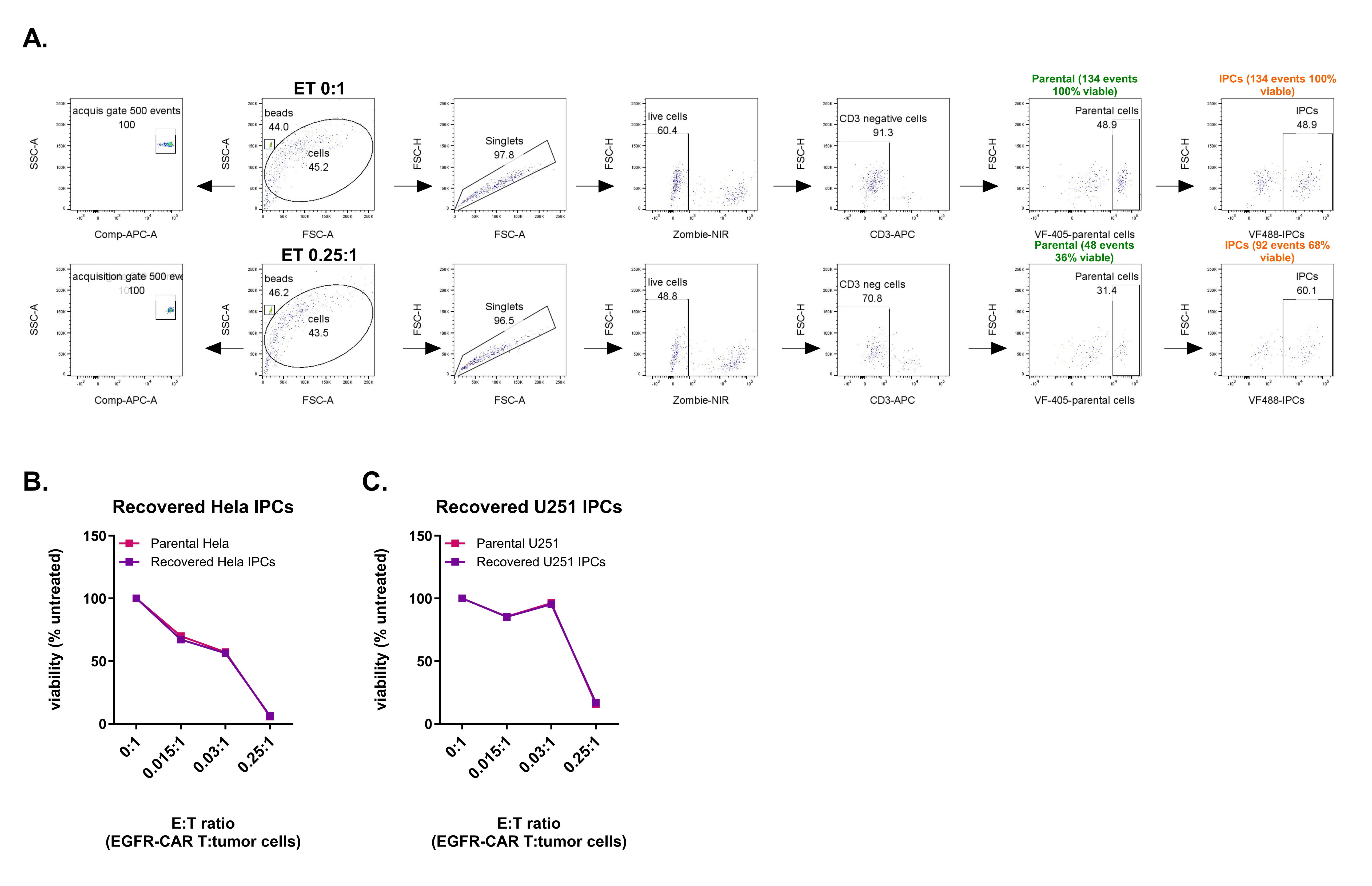
